# IgG-Bridging–Seeded Synergistic Aggregation of SARS-CoV-2 Spikes Underlies Potent Neutralization by A Low-Affinity Antibody

**DOI:** 10.1101/2025.11.30.691454

**Authors:** Niannian Lv, Peng Chen, Xiaobin Dai, Hu Xu, Ziheng Li, Zelin Shan, Jinqian Li, Fenglin Guo, Yuanfang Chen, Jiayi Li, Yiqian Huang, Guizhi Dong, Yifan Jiang, Liang Chen, Xuanyu Nan, Hanjun Zhao, Kang Zhang, Shilong Fan, Yuanchen Dong, Dongsheng Liu, Xinquan Wang, Deli Huang, Xiaojing Pan, Chunying Chen, Zhihua Liu, Li-Tang Yan, Qi Zhang, Lin Qi Zhang, Yuliang Zhao, Yuhe Renee Yang

## Abstract

Mechanistic studies of viral neutralization typically prioritize high-affinity antibodies, relegating low-affinity binders to the sidelines. We report P5-1C8, a Class 1 SARS-CoV-2 antibody that exemplifies this underexplored “low-affinity yet high-potency” phenotype, retaining strong neutralization of Omicron JN.1 despite markedly weakened trimer binding (K_D_ = 225 nM; IC_50_ = 0.06 nM). Structural and biophysical analyses reveal that P5-1C8 engages WT and BA.1 spikes through canonical intra-spike bivalency, but with JN.1 it induces aggregation. Using virion-like nanoparticles displaying multiple spikes, we show that IgG remains bound with no detectable dissociation and triggers pronounced aggregation. Coarse-grained molecular dynamics delineate the stepwise pathway in which weak IgG-spike contacts seed aggregation via transient inter-spike bridging. Together, these findings establish the first mechanistic framework demonstrating how weak-binding antibodies can nonetheless achieve potent neutralization through higher-order aggregation, thereby expanding the conceptual landscape of antibody function and opening new directions for antibody evaluation and design.

**Graphical Abstract:** Low-affinity antibodies are frequently disregarded in discovery pipelines. This work reports P5-1C8, a Class 1 SARS-CoV-2 antibody with weak trimer binding (K_D_-to-IC_50_ > 3,700-fold) yet potent neutralization of Omicron JN.1. Structural, biophysical, functional and coarse-grained simulations collectively demonstrate that transient inter-spike IgG bridging seeds higher-order aggregation, which in turn drives neutralization and provides a mechanistic framework.

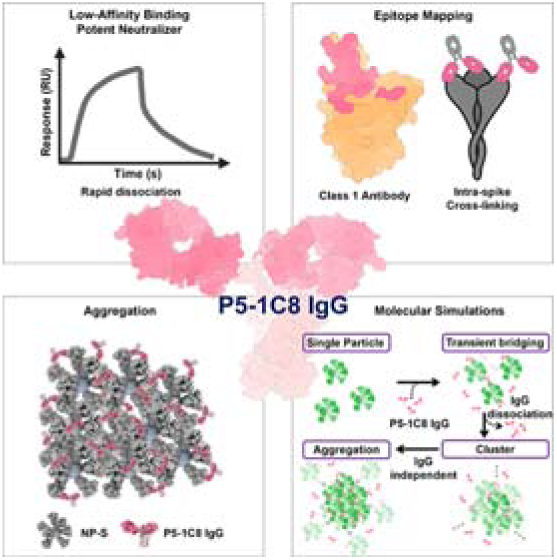

## 1. Introduction

Neutralizing antibodies are central to humoral immunity against viral infections.^[1]^ They employ diverse mechanisms, ranging from blocking viral attachment by competing with receptor binding,^[2]^ inducing spike conformational changes,^[3]^ or promoting virion aggregation;^[4]^ to interfering with membrane fusion,^[5]^ endosomal cleavage, or receptor engagement after attachment;^[6]^ to restricting viral spread by suppressing progeny release^[7]^ or preventing direct cell-to-cell transmission.^[8]^ Despite their mechanistic diversity, all these processes converge on a common initiating event—binding. In the case of severe acute respiratory syndrome-coronavirus-2 (SARS-CoV-2), numerous studies have focused on antibodies with high-affinity and potent neutralization,^[9]^ with general expectation that binding affinity—or more accurately, avidity—correlates with neutralization.^[10]^ Using a combined dataset of equilibrium dissociation constants (K_D_) and neutralization potencies (IC_50_) for coronavirus-related antibodies, compiled from the Ab-CoV database (reported up to 2022; https://web.iitm.ac.in/bioinfo2/ab-cov/covabseq/),^[11]^ and supplemented with additional data published after 2022, we noted that affinity and avidity measurements are almost exclusively derived from binding to monomeric receptor-binding domains (RBDs), rather than to trimeric spike proteins. While this approach enables the distinction between Fab-RBD and IgG-RBD interactions, which often revealing affinity or avidity differences ranging from several-fold to several hundred-fold. It overlooks key structural features of the native spike, such as dynamic “up” and “down” RBD conformations,^[12]^ as well as critical factors of bivalent IgG binding, including Fab approach angles,^[13]^ post-binding stoichiometry,^[14]^ and multivalent binding In contrast to the vast majority of studies that rely on monomeric domains, only a handful of reports to date, have examined the binding of Fabs and full-length IgGs to the intact spike trimer.^[13, 16]^ From these limited trimer-based studies, a consistent observation has emerged: high-affinity Fabs (sub-nanomolar) confer potent neutralization in both Fab and IgG formats, whereas moderate-affinity Fabs (tens of nanomolar) lose neutralizing activity in Fab fragment form but retain potency as full-length IgGs due to bivalent engagement.^[13]^ Rare exceptions, such as COVA1-03^[9b]^ and COVA1-25,^[9b]^ display undetectable apparent avidity yet maintain strong neutralizing activity. These antibodies have received little mechanistic investigation, likely due to their rarity, the prevailing emphasis on high-affinity candidates in antibody discovery pipelines,^[9]^ and a methodological disconnect where binding is typically measured using monomeric RBDs or Fab fragments, whereas neutralization is measured using full-length IgGs against pseudovirus or live virus.^[17]^

To elucidate the mechanism of antibody-mediated neutralization, structural biology is widely employed to map epitopes, define binding interactions, and reveal the modes of neutralization.^[18]^ Most studies to date have focused on Fab-RBD complexes using X-ray crystallography^[14c, 19]^ or Fab-spike complexes using cryo-electron microscopy (cryoLJEM),^[20]^ approaches that provide high-resolution insight into epitope targeting but fail to capture avidity effects and Fc-mediated contributions.^[21]^ In contrast, analyses of full-length W328-6H2 IgG have revealed distinct, variant-specific higher-order architectures that directly affect its neutralizing capability.^[16a]^ Such higher-order IgG-spike assemblies can reveal previously inaccessible dimensions of antibody function that are not captured by conventional structural approaches. For instance, while antibody P17 was previously known to neutralize SARS-CoV-2 by blocking ACE2 binding and inhibiting membrane fusion,^[22]^ and S309 was recognized for targeting the conserved N343 glycan and exhibiting enhanced Fc-dependent effector functions.^[16b]^ However, visualization of full-length IgG interacting with native spike trimers on the viral surface has uncovered previously unrecognized mechanisms. For P17, cryo-electron tomography revealed that bivalent IgG assembles spike trimers into linear multimers, likely generating mechanical tension that induces S1 shedding—an effect not apparent from Fab-based or isolated trimer studies.^[23]^ Similarly, for S309, higher-order pentameric and hexameric lattice structures were observed,^[23]^ providing a structural basis for enhanced complement activation,^[24]^ antibody-dependent cell cytotoxicity (ADCC), antibody-dependent cellular phagocytosis (ADCP), and FcγRIIa signaling.^[16b, 23]^ Together, these findings demonstrate that Fab-based structures alone cannot capture the full scope of antibody function. A comprehensive understanding of neutralization mechanisms requires multiscale structural analyses—from Fab and full-length IgG, and from RBD to spike trimer to virion—combined with diverse biophysical and functional assays.

In this work, we describe P5-1C8, a broadly neutralizing antibody isolated from a convalescent individual infected with the ancestral SARS-CoV-2 strain. Despite exhibiting markedly reduced binding avidity to the Omicron subvariant JN.1 (K_D_ = 225 nM), P5-1C8 retains potent neutralization activity (IC_50_ = 0.06 nM), resulting in a >3,700-fold K_D_-to-IC_50_ disparity. XLJray crystallography mapped its epitope to the receptorLJbinding motif (RBM), confirming its ClassLJ1 category, which almost invariably employ the canonical intra-spike bivalent binding mode.^[13]^ NegativeLJstain and single-particle cryo-EM revealed that P5-1C8 adopts the standard intra-spike bivalent geometry on both wild-type (WT) and Omicron BA.1 spikes, in line with prior structures.^[13]^ In contrast, no such binding mode was detected with JN.1; instead, P5-1C8 promoted extensive aggregation. This behavior was corroborated using virion-like nanoparticles displaying multiple spikes, where multivalent presentation abolished measurable IgG dissociation and drove large-scale aggregation. Coarse-grained molecular dynamics further delineated the process: on WT nanoparticles, P5-1C8 engaged in high-avidity intra- and inter-spike binding mode, whereas weaker binding to JN.1 spikes seeded transient assemblies which initiate the stable aggregation. Together, these results reveal that low-affinity antibodies—even those ineffective in monovalent assays—can exert potent neutralization by a previously underappreciated mechanism of higher-order assembly, and expands the conceptual farmwork of antibody function and underscores the need to systematically include such antibodies in mechanistic studies, therapeutic discovery, and vaccine evaluation.

## 2. Results

### 2.1. Neutralization potency, binding affinity and epitope mapping of antibody P5-1C8

As detailed in our previous published work,^[19a]^ we isolated a broadly neutralizing antibody, P5-1C8, from a convalescent individual (P5) following ancestral SARS-CoV-2 infection. P5-1C8 demonstrated exceptional neutralizing potency against wild-type (WT) SARS-CoV-2 virus (IC_50_ = 0.014 μg/mL), and maintained strong neutralizing activity across a panel of major SARS-CoV-2 variants, including Beta, Delta, BA.1, BA.2.75, BA.5, BF.7, BQ.1, BQ.1.1, XBB, XBB.1, and JN.1, with IC_50_ values ranging from 0.003 to 0.042 μg/mL (**Figure 1**a-c; Figure S1, Supporting Information). To investigate the mechanistic basis underlying the broad and potent neutralizing activity of P5-1C8, we first characterized the binding kinetics of full-length P5-1C8 IgG using surface plasmon resonance (SPR), with His-tagged trimeric spike proteins immobilized on nitrilotriacetic acid (NTA) sensor chips. P5-1C8 displayed extremely high avidity for WT spike trimer (K_D_ < 1 pM, Figure 1d), and retained strong binding for BA.1 spike trimer (K_D_ ≈ 8.7 nM, Figure 1e). Remarkably, however, its binding to JN.1 spike trimer was substantially reduced, with a K_D_ of approximately 225 nM—nearly three orders of magnitude weaker than BA.1 and over five orders weaker than WT (Figure 1f). While the association rate constant (K_a_) remained comparable across WT, BA.1, and JN.1 spikes, the dissociation rate constant (K_d_) for JN.1 was elevated by approximately six orders of magnitude compared to WT spike. Despite this markedly reduced binding avidity, P5-1C8 retained potent neutralization against JN.1, with an IC_50_ of 0.009 μg/mL (Figure 1c). The striking disconnect between binding avidity and neutralization potency—a ∼3750-fold divergence between K_D_ and IC_50_—represents a profound decoupling of antigen recognition from antiviral efficacy. This observation stands in sharp contrast to prevailing models of antibody-mediated neutralization, in which affinity or avidity is generally predictive of functional potency (Table S1, Supporting Information).

**Figure 1.**
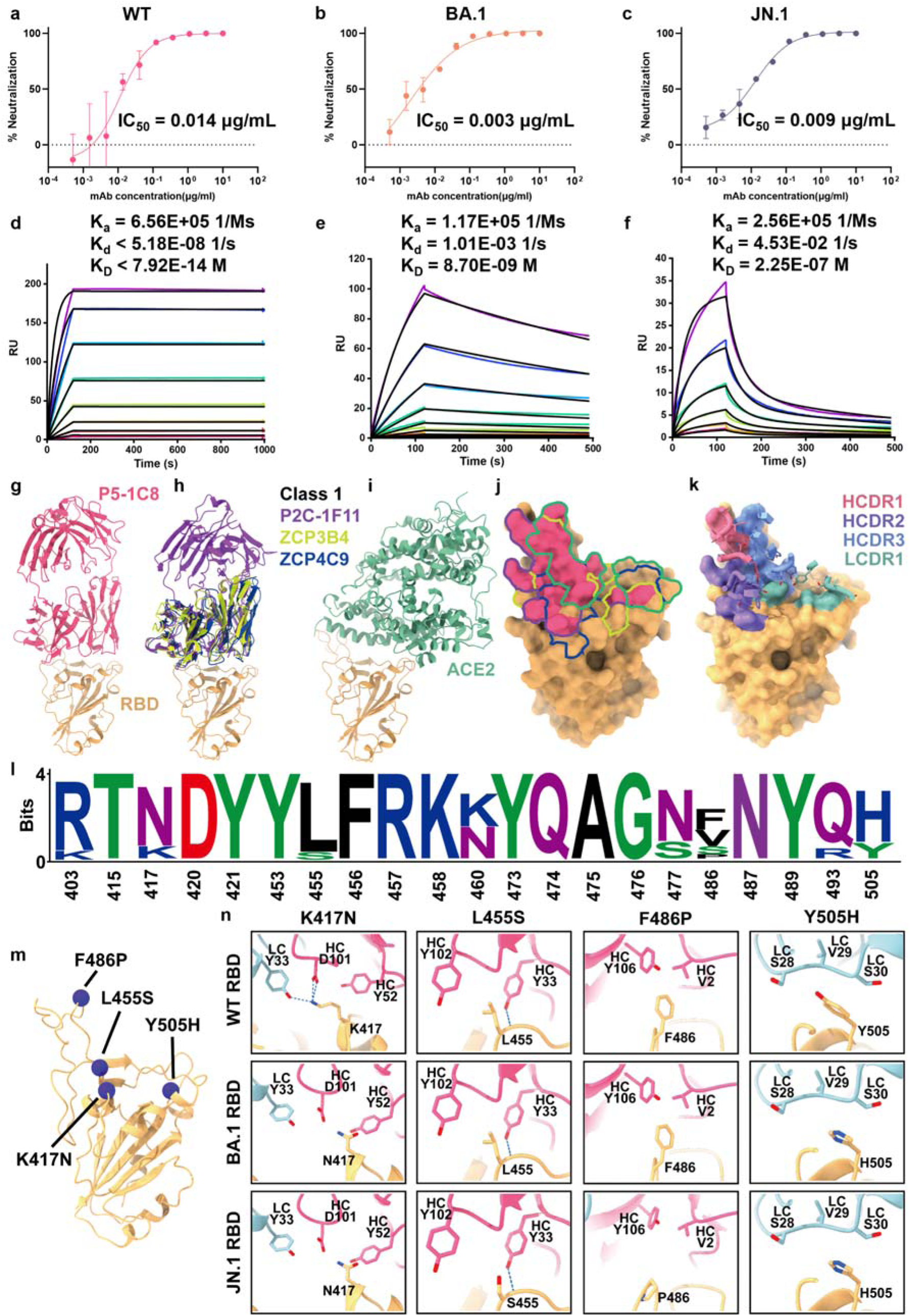
Neutralization potency, binding affinity and epitope mapping of antibody P5-1C8. a-c) Neutralization c0075rves of P5-1C8 IgG against SARS-CoV-2 WT (a), BA.1 (b), and JN.1 (c) variants. Data was obtained from two independent experiments, each performed with two technical replicates. d-f) Binding kinetics between P5-1C8 IgG and the spike trimer of SARS-CoV-2 WT (d), BA.1 (e), and JN.1 (f), measured by surface plasmon resonance (SPR). Spike trimers were immobilized on a nitrilotriacetic acid (NTA) sensor chip, and serial dilutions of P5-1C8 IgG were flowed through the system. Colored lines represent experimentally measured sensorgrams. Black lines show the best-fit curves based on experimental data. The calculated association rate (K_a_), dissociation rate (K_d_), and equilibrium dissociation constant (K_D_) for each antibody-spike pair are indicated. The dissociation rate constant of P5-1C8 IgG for WT spike should be interpreted with caution, as it near the detection limit of instrument. All results were confirmed in three independent experiments. g) Overall structure of the complex between SARS-CoV-2 WT RBD (soft orange) and P5-1C8 Fab (pink). h) Superimposition of three Fab-RBD complexes: P2C-1F11 (dark violet), ZCP3B4 (moderate yellow), and ZCP4C9 (blue). i) Crystal structure of WT RBD (soft orange) in complex with ACE2 (green), panels g-i show the spatial relationships of all four Fabs relative to the ACE2 binding site. j) Binding footprints of the four Fabs and ACE2 on the SARS-CoV-2 RBD. Pink, dark violet, moderate yellow, blue, and green denote the footprints of P5-1C8 Fab, P2C-1F11 Fab, ZCP3B4 Fab, ZCP4C9 Fab, and ACE2, respectively. k) Footprint of RBD-contacting residues on antibody CDR loops, with key amino acids in HCDR1-3 and LCDR1. CDR positions are annotated according to IMGT numbering. l) Sequence conservation analysis on residues bound by P5-1C8. These logo plots show the conservation of P5-1C8 epitopes from SARS-CoV-2 WT, Beta, Delta, and Omicron (BA.1, BA.2.75, BA.5, BF.7, BQ.1, BQ.1.1, XBB, XBB.1, and JN.1) variants. m) Structure of the SARS-CoV-2 WT RBD, with key P5-1C8 Fab-binding residues highlighted as violet spheres. n) The interactions between P5-1C8 Fab and non-conserved residues in the RBD are affected by strain-specific mutations during viral evolution. Shown are the structural comparisons of P5-1C8 Fab interactions with WT RBD (upper panel) and Omicron BA.1 RBD (middle panel), and JN.1 RBD (lower panel) at positions K417N, L455S, F486P, and Y505H. Contacting residues are depicted as sticks, and hydrogen bonds or salt bridges are indicated by dashed lines.

Given the striking disconnect between binding avidity and neutralization potency, we next sought to define the structural basis of P5-1C8 recognition through high-resolution epitope mapping. To this end, we determined the crystal structure of the P5-1C8 Fab in complex with the wild-type SARS-CoV-2 RBD at 2.39-Å resolution (Figure 1g; Figure S2, Supporting Information). Structural alignment with three previously reported Fab-RBD complexes revealed that P5-1C8 shares a nearly identical binding pose with P2C-1F11,^[19d]^ ZCP3B4,^[20d]^ and ZCP4C9,^[20d]^ targeting the top face of RBD, mimicking the binding mode of ACE2 (Figure 1h,i).^[19d]^ All four antibodies adopt a similar footprint as well as orientation characteristic of class 1 RBD-targeting antibodies (Figure 1j; Figure S3, Supporting Information). The P5-1C8 paratope comprises 21 residues, with 17 contributed by the heavy chain (6 from HCDR1, 5 from HCDR2, 5 from HCDR3) and 4 from the light chain (all from LCDR1), burying a total surface area of ∼1003.7 Å² (Figure 1k). The interface includes 15 hydrogen bonds and two salt bridges (Table S4, Supporting Information), and exhibits substantial epitope overlap with P2C-1F11 (21 shared residues), ZCP3B4 (18), ZCP4C9 (19), and ACE2 (10) (Figure S3, Supporting Information), confirming that P5-1C8 targets a conserved class 1 epitope. To assess conservation, we performed multiple sequence alignments across variants of concern (VOCs; Figure S4, Supporting Information), revealing that 8 of the 21 contact residues are mutated in at least one variant (Figure 1l). Structural analysis of the P5-1C8-RBD interface in WT, BA.1, and JN.1 showed that four substitutions (R403K, N460K, S477N, Q493R) caused minimal disruption (Figure S5, Supporting Information), whereas K417N, L455S, F486P, and Y505H significantly altered key interactions (Figure 1m,n). For example, K417N abolishes cation-π and salt-bridge interactions;^[25]^ L455S increases flexibility and weakens hydrophobic contacts;^[26]^ F486P disrupts π-π stacking and destabilizes local structure;^[27]^ and Y505H impairs both hydrophobic and aromatic interactions.^[28]^ These changes collectively contribute to the observed stepwise reduction in binding avidity from WT to BA.1 to JN.1, as reflected in SPR measurements. Although structural and kinetic analyses align in defining the epitope and affinity/avidity landscape, they cannot account for the potent neutralizing activity observed against JN.1, suggesting that P5-1C8 may engage in an alternative or auxiliary neutralization mechanism beyond conventional receptor blockade.

### 2.2. Multivalent binding of full-length P5-1C8 IgG to WT, BA.1 and JN.1 spike trimers

To dissect how P5-1C8 engages SARS-CoV-2 spike trimers from the structural perspective, we employed negative-stain electron microscopy (ns-EM) to characterize the global structure of P5-1C8 both in full-length IgG and Fab fragments in complex with WT, BA.1, and JN.1 spikes (**Figure 2**a,b; Figure S6, Supporting Information). We grouped the resulting two-dimensional (2D) averages based on the number of Fab molecules bound per spike trimer and quantified particle distributions within each class to estimate binding occupancy (Figure 2c), which was subsequently correlated with SPR-derived binding affinities (Figure 2d; Figure S7, Supporting Information). Upon binding to WT spike, both full-length IgG and Fab forms of P5-1C8 exhibited high occupancy, engaging up to three RBDs per trimer (Figure 2a,b). Over 95% of IgG-spike particles showed 2-3 bound Fabs in 2D class averages, with full occupancy confirmed by 3D reconstruction; Fab displayed >93% occupancy with 3 Fabs per trimer (Figure 2a-c). On average, IgG bound 1.5 molecules per trimer, while Fab saturated all three RBDs. Both forms exhibited picomolar-range KD values (Figure 2d) and potent neutralization, with IC_50_ values of 0.014 μg/mL for IgG (Figure 1a) and 0.018 μg/mL for Fab (Figure S7a, Supporting Information), consistent with previously reported high-affinity class 1 antibodies.^[13]^

**Figure 2.**
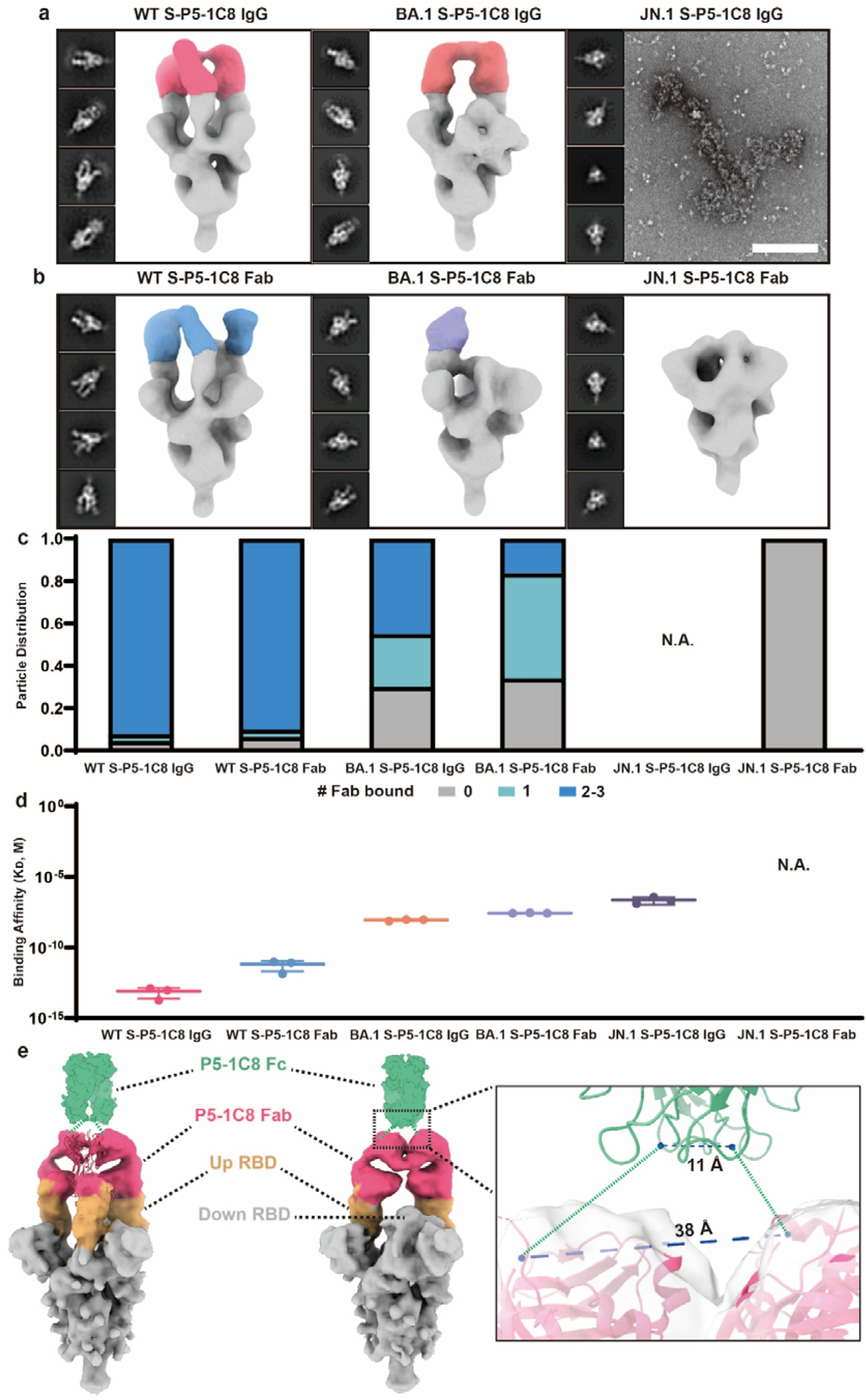
Multivalent binding analysis of antibody P5-1C8 to WT, BA1 and JN1 spike trimers. a) Representative two-dimensional (2D) class averages and side views of composite 3D reconstructions from ns-EM of P5-1C8 IgG in complex with spike proteins from SARS-CoV-2 WT (pink), BA.1(red) and JN.1, at a 3:1 molar ratio (IgG/Spike). In the JN1 S-P5-1C8 IgG complex, particles do not form well-defined complexes but instead aggregate upon binding. Scale bar: 200 nm. b) Representative 2D class averages and side views of composite 3D reconstructions from ns-EM of P5-1C8 Fab in complex with spike proteins from SARS-CoV-2 WT (blue), BA.1(purple) and JN.1, at a 3:1 molar ratio (Fab/Protomer). No binding is observed in the JN.1 S-P5-1C8 Fab complex. c) Semi-quantitative epitope occupancy analysis derived from ns-EM shown in a and b, showing the proportion of spike trimers bound by 0, 1, 2-3 Fabs (gray, light cyan, and blue, respectively). Data corresponds to the S-P5-1C8 IgG (a) and S-P5-1C8 Fab (b) complexes for SARS-CoV-2 WT, BA.1, and JN.1, as indicated for each participant on the x-axis. d) Binding affinity values obtained from SPR assays of P5-1C8 IgG and Fab against SARS-CoV-2 WT, BA.1, and JN.1 spikes. Binding of P5-1C8 Fab to JN.1 spike was not detectable. N.A., not available. All experiments were performed independently in triplicate. e) Cryo-EM map of the SARS-CoV-2 WT spike in complex with P5-1C8 IgG. The reconstruction reveals two binding stoichiometries: a symmetric complex (1.5 IgG bound) and an asymmetric complex (one IgG bound). Up RBDs, down RBD, and P5-1C8 Fabs are shown in yellow, grey, and pink, respectively. The Fc region of P5-1C8, shown in green, was modeled using AlphaFold 3 predictions based on the P5-1C8 Fc sequence. The absence of Fc density in the WT S-P5-1C8 IgG reconstruction is likely due to intrinsic hinge-mediated flexibility of the IgG molecule.^[16c]^ The zoomed-in view on the right highlights the structural details within the black dashed box. The two CH1 C-termini of the Fabs are separated by 38 Å, whereas the N-termini of the Fc heavy chains are spaced 11 Å apart.

In contrast to WT, P5-1C8 exhibited reduced binding to the BA.1 spike. Ns-EM analysis showed a decrease in IgG occupancy to ∼70%, with most particles bound by two Fabs and 3D reconstructions confirming bivalent engagement (Figure 2a,c). The Fab form displayed further reduced binding (∼66%), with the majority of particles showing only one Fab per trimer (Figure 2b,c) and ∼16% exhibiting two (Figure 2c; Figure S8, Supporting Information). Notably, full-length IgG retained bivalent engagement while the third RBD site bound with monovalent Fab was no longer detectable, indicating that avidity allows IgG to compensate for reduced intrinsic affinity. Consistently, Fab exhibited markedly reduced binding and a ∼1,140-fold drop in neutralization potency (IC_50_ = 3.422 μg/mL; Figure S7b, Supporting Information), while the IgG form maintained high avidity binding and potent neutralization.

To further resolve the precise binding geometry, we determined single particle cryo-EM structures of the P5-1C8 IgG bound to SARS-CoV-2 WT spike trimers (Figure S9, Supporting Information). We identified two distinct populations of WT S-P5-1C8 IgG complexes (Figure 2e). In state 1, the spike trimer adopts a symmetric conformation with all three RBDs in the “up” position. Each trimer accommodates approximately 1.5 IgGs, with one IgG binding two RBDs and the third RBD bound by a separate IgG, potentially bridging adjacent trimers. In state 2, an asymmetric trimer displays two “up” RBDs, both engaged by two Fabs from the same IgG. By docking the crystal structure of RBD-P5-1C8 Fab complex (PDB 9K6J) into the EM densities, we measured a distance of 38 Å between Pro217 residues of the heavy chains—well within the ∼65 Å span allowed by the IgG hinge^[20a, 23]^—thereby confirming that the two Fab densities originated from a single IgG molecule and supporting the model of intra-spike bivalent binding (Figure 2e).

Unexpectedly, P5-1C8 IgG bound the JN.1 spike in an atypical manner, with particles showing either pronounced trimer aggregation or no detectable IgG-spike complexes (Figure 2a; Figure S6, Supporting Information). Negative-stain 2D class averages of well-dispersed particles confirmed only spike classes without discernible IgG density, indicating the lack of stable complex formation (Figure S10, Supporting Information). To further probe this interaction, we incubated P5-1C8 IgG with JN.1 spike trimers at an IgG-to-spike molar ratio of 3 (∼1.7 μM IgG to ∼0.6 μM spike) for durations ranging from 5 mins to 4 h. In all micrographs, dispersed spike trimers showed no detectable evidence of IgG engagement (Figure S11, Supporting Information), suggesting that binding occurred exclusively within aggregated clusters. At ∼28,000-fold lower IgG concentration (0.06 nM, approximating the IC_50_), dispersed particles remained unbound, with only sparse aggregates observed; further reduction to one-tenth of the IC_50_ abolished aggregation entirely (Figure S12, Supporting Information). Across all tested conditions, no stable complex between P5-1C8 IgG and JN.1 spike are captured. The Fab form showed no binding, no aggregation, and no neutralizing activity (IC_50_ > 10 μg/mL, Figure 2b-d; Figure S7c,f, Supporting Information). Unlike the strong binding observed with P5-1C8 IgG to WT spike, JN.1 spike showed no detectable binding with monovalent Fab, and moderate binding to full-length IgG (K_D_ = 225 nM) and induced aggregation, suggesting that IgG bivalency engages multiple spike trimers.

### 2.3. Multivalent binding mode of full-length P5-1C8 IgG with NP-S nanoparticles

To investigate how full-length P5-1C8 IgG engages multiple trimeric spikes, we used nanoparticles (kindly provided by collaborator) coated with spike proteins (NP-S) to mimic the multivalent display on intact virions (**Figure 3**a). We first measured this interaction between P5-1C8 IgG and NP-S nanoparticles by SPR (Figure 3b,c). To directly compare the binding behavior of soluble spike and NP-S, P5-1C8 IgGs were immobilized on Protein A-coated CM5 sensor chips, followed by injection of either soluble spike (Figure 3b) or NP-S nanoparticles (Figure 3c). P5-1C8 IgG exhibited potent binding avidity, with comparable association rates for soluble WT S (4.73 × 10^6^ Ms^-1^) and NP-WT S (1.63 × 10^6^ Ms^-1^), and extremely slow dissociation rates near the detection limits of instrument (Figure 3b,c). In contrast, association rates were also similar between soluble and NP-displayed JN.1 spikes, but their dissociation rates diverged sharply as the soluble form dissociated rapidly (1.00 × 10^-2^ s^-1^; Figure 3b), whereas NP-presented form exhibited dramatically slower off-rates (4.76 × 10^-6^ s^-1^; Figure 3c), indicating enhanced stability driven by multivalent interactions. Similarly, cell surface staining assay also analyzes multivalent binding of P5-1C8 IgG to multiple spikes. By incubating IgG with HEK293T cells expressing various spike proteins from WT to JN.1 SARS-CoV-2 variants, revealing comparable binding across all variants with only a modest 1.5-fold reduction for JN.1 compared to WT (Figure S13, Supporting Information). Compared with the free spike system, systems presenting multiple trimeric spikes, either on nanoparticles or cell surfaces, enable stable binding of full-length IgG.

**Figure 3.**
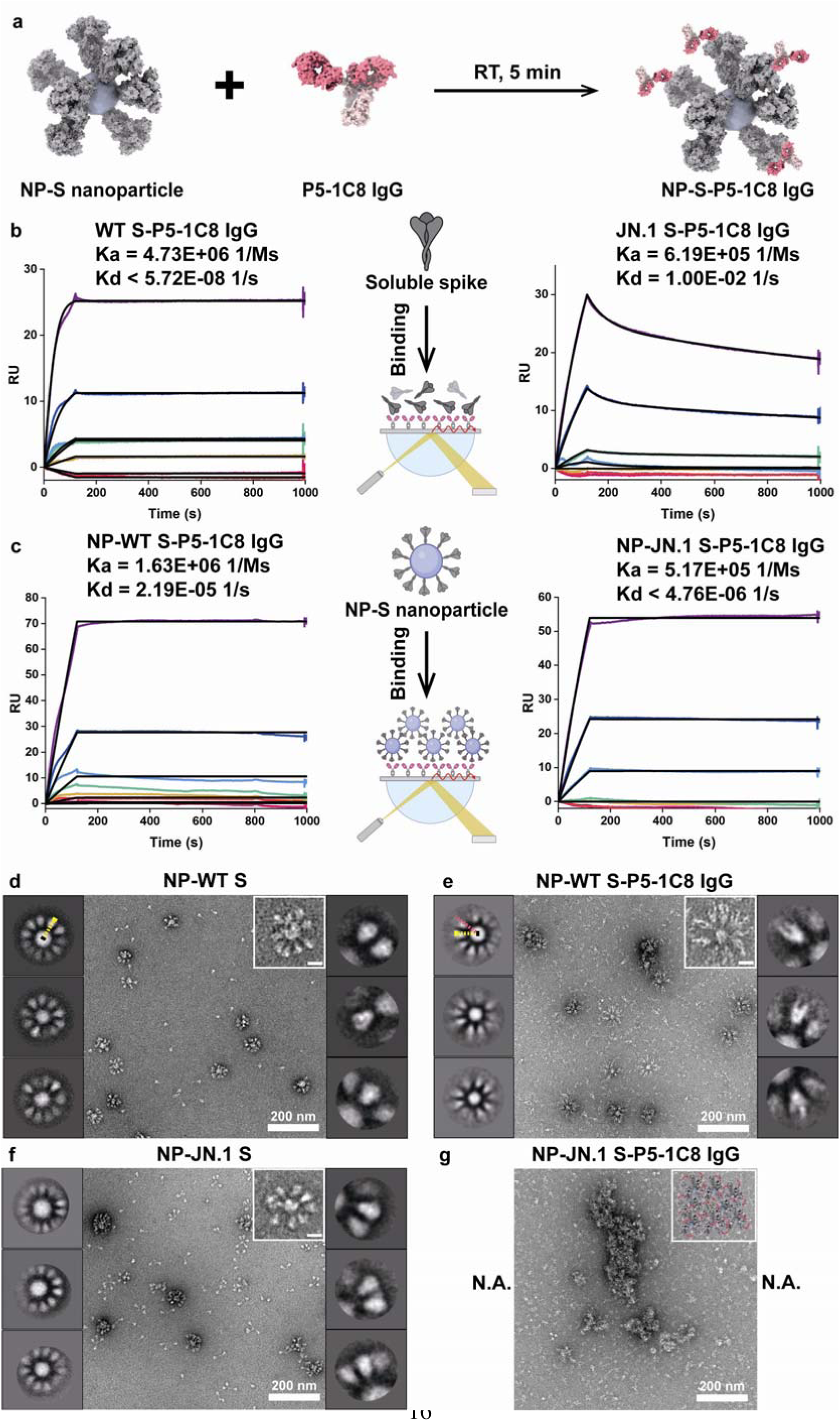
Multivalent binding mode of full-length P5LJ1C8 IgG with NP-S nanoparticles. a) Schematic illustration of P5-1C8 IgG binding to NP-S nanoparticle. b,c) SPR analysis of P5-1C8 IgG binding to either soluble spike trimers (b) or NP-S nanoparticles (c) from WT (left) and JN.1 (right). Protein A was immobilized on a CM5 sensor chip to capture P5-1C8 IgG, and the serial dilutions of either solution spike trimers or NP-S nanoparticles were flowed through the system. Colored lines represent the raw sensorgrams. Black lines show the best-fit curves based on experimental data. The calculated association rate (K_a_) and dissociation rate (K_d_) are shown. All results were performed in three independent experiments. The schematic illustrations in the middle panels of (b) and (c) are created with BioRender.com. d,e) Ns-EM images (middle) and 2D class averages (left and right panels) of NP-WT S nanoparticles before (d) and after (e) incubation with P5-1C8 IgG. Scale bar: 200 nm. Inset (upper right) shows zoomed-in view of a representative nanoparticle. Scale bar: 20 nm. In d (left panel), an auxiliary radius was defined from the NP-WT S nanoparticle center to the spike tip, yielding ∼30.9 nm. When the same reference radius was applied to e (left panel), the pink line extended ∼7.1 nm beyond it, with this offset corresponding to bound IgG density. f,g) Ns-EM images (middle) and 2D class averages (left and right panels) of NP-JN.1 S nanoparticles before (f) and after (g) addition of P5-1C8 IgG. Scale bar: 200 nm. The upper right insets show a zoomed-in view of a representative nanoparticle (left) and a schematic illustration of antibody-induced aggregation (right). The left and right panels of the ns-EM micrographs in Figure d to f show NP-S nanoparticles or NP-S-IgG complexes and spike or spike-IgG complexes on nanoparticle, respectively, illustrating representative 2D class averages.

We next performed negative-stain EM to examine the binding mode of P5-1C8 IgG to NP-S nanoparticles (Figure 3d-g). Comparable antibody and spike concentrations were used across NP and free spike systems (∼1.4 μM IgG to ∼0.5 μM spike trimer displayed on nanoparticles). Prior to antibody addition, negative-stain micrographs and 2D class averages showed uniformly distributed spikes projecting outward from NP surface (Figure 3d,f). We defined an auxiliary radius extending from the NP-WT S nanoparticle center to the spike tip, yielding ∼30.9 nm (Figure 3d, left panel). After IgG incubation, most nanoparticles remained monodisperse. Applying the same reference radius to NP-WT S-P5-1C8 IgG complexes revealed an additional ∼7.1 nm density beyond the reference, attributable to bound IgG (Figure 3e, left panel). Zoomed-in views and 2D class averages of selected spikes on nanoparticles further confirmed bivalent binding to NP-displayed spikes (Figure 3d,e, right panel). Conversely, NP-JN.1 S exhibited aggregation (Figure 3g). When incubation time was extended to 10 or 30 mins, larger and more abundant aggregates were observed in each micrograph (Figure S14, Supporting Information), whereas incubation with monovalent P5-1C8 Fab resulted in neither binding nor aggregation (Figure S15, Supporting Information), indicating that the bivalency of full-length IgG is required for cluster formation, similar to the trend seen in free-solution experiments (Figure S11, Supporting Information). To assess antibody-induced aggregation under native, virion-like conditions, Omicron JN.1 pseudoviruses were incubated with P5-1C8 IgG and analyzed by negative-stain EM. Aggregation of virus particles was observed (Figure S16, Supporting Information), confirming that IgG promotes JN.1 particle clustering. Overall, enhanced IgG binding were observed with spikes displayed on nanoparticles compared with free spikes, potentially due to multivalent binding to adjacent WT spikes on nanoparticles, while JN.1 revealed only particle aggregation, independent of incubation time, and the underlying process remains unclear.

### 2.4. Molecular dynamics simulations reveal aggregation mechanism of P5-1C8 IgG

To investigate how weakly binding P5-1C8 IgG drives aggregation of JN.1 spike and NP-JN.1 S, we employed coarse-grained (CG) molecular dynamics simulations,^[29]^ which capture both local binding events and global aggregation. CG models of WT and JN.1 spike trimers and full-length IgG were constructed to preserve domain architecture and flexibility for bivalent engagement. Specifically, spike trimers were partitioned into four structural segments: residues 1-527 (NTD and RBD), 528-833 (CTD1, CTD2, and FP), 834-1162 (HR1, CH, and CD), and 1163-1240 (HR2 and foldon);^[30]^ and full-length P5-1C8 IgG was modeled with two Fab fragments, a flexible hinge, and an Fc domain (**Figure 4**a).^[31]^ Simulations were configured with eight spike trimers displayed on 15 nm nanoparticles, as estimated from prior ns-EM data, and parameterized using experimental KD values and concentrations (1.4 μM IgG and ∼0.5 μM spike trimer). Each simulation box (1500×1500×1500 nm^3^) contained 125 NP-S particles, and trajectories were run for up to 15 μs. Upon IgG binding, NP-WT S remained monodisperse throughout the trajectory (Figure 4b, top; Movie S1, Supporting Information), whereas NP-JN.1 S exhibited progressive clustering (Figure 4b, bottom; Movie S2, Supporting Information). To quantify these dynamics, we monitored average cluster occupancy, cluster counts, and the populations of free versus bound IgG and nanoparticles (Figure 4c-f). Clusters were defined as assemblies with center-to-center separations of less than one nanoparticle diameter. For NP-WT S, the number of IgG-bound nanoparticles rapidly plateaued, while the number of nanoparticle-bound IgG continued to increase, indicating spatially uniform binding across all nanoparticles in the simulation volume and reaching saturation (Figure 4c). This yielded monodisperse NP-WT S/IgG complexes without detectable clustering (Figure 4d,e). In contrast, NP-JN.1 S aggregated through a two-step process. During the initial phase, rapid IgG binding coincided with a synchronous increase in the number of IgG-bound nanoparticles (Figure 4f) and in cluster counts (Figure 4d). These clusters remained small, with average occupancies below three (Figure 4e), and IgG incorporation peaked at ∼14.2% (Figure 4f). In the second phase, both bound IgG and IgG-bound nanoparticles declined, while clusters expanded, reflected by fewer total clusters and a steady increase in average occupancy, reaching equilibrium at ∼11 nanoparticles per cluster (Figure 4d-f). Notably, almost all nanoparticles were incorporated into clusters, whereas only ∼2.1% of IgG remained bound (Figure 4f), indicating that IgG primarily drives step-1 nucleation but is dispensable for step-2 growth into larger aggregates. Cryo-EM analysis at the same IgG concentration (1.4 μM) corroborated the simulation results (Figure 4g,h). Prior to IgG addition, NP-S were uniformly distributed. After IgG incubation, NP-WT S remained as monodisperse complexes (Figure 4g), while NP-JN.1 S exhibited amorphous aggregates that predominated across the fields of view (Figure 4h).

**Figure 4.**
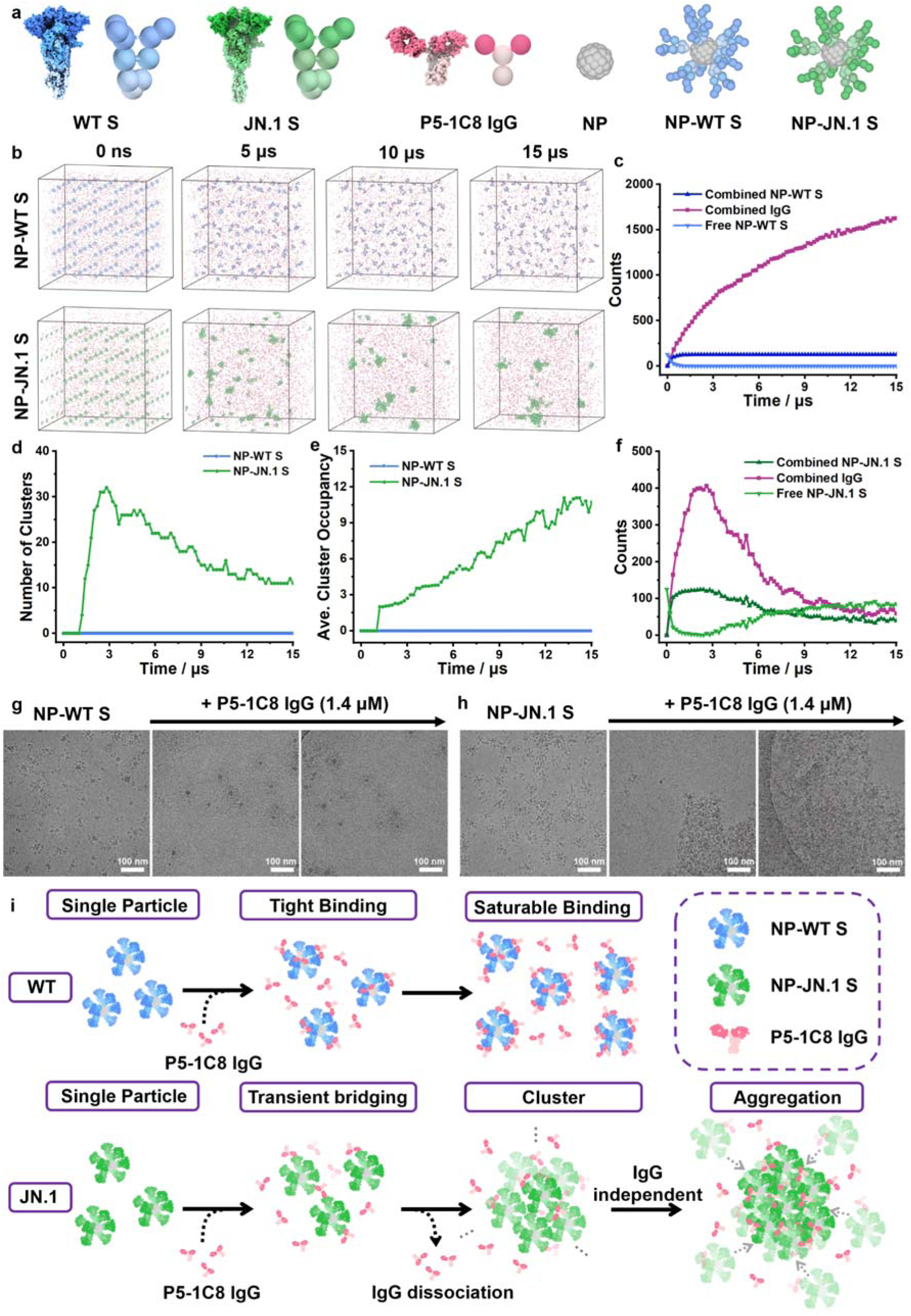
Coarse-grained molecular dynamics simulations of P5-1C8 IgG interactions with WT and JN.1 spikes displayed on nanoparticles. a) Coarse-grained bead representation of spike proteins, IgG molecule and nanoparticles. WT S is shown in blue, JN.1 S in green, P5-1C8 IgG in pink, and nanoparticles in grey. b) Representative snapshots of the binding process between P5-1C8 IgG and either NP-WT S (top) and NP-JN.1 S (bottom). Simulations were conducted with 1.4 μM IgG and ∼0.5 μM spike trimer displayed on nanoparticles, within a cubic box of 1500×1500×1500 nm^3^ containing 125 NP-S particles, with trajectories were run up to 15 μs. c) Temporal evolution of the number of free or bound P5-1C8 IgG molecules and NP-WT S nanoparticles within the simulation box. d,e) Temporal evolution of (d) cluster number and (e) average cluster occupancy in NP-WT S (blue) or NP-JN.1 S (green) systems during simulations with P5-1C8 IgG. f) Temporal evolution of the number of free or bound P5-1C8 IgG molecules and NP-JN.1 S nanoparticles within the simulation box. g,h) Representative cryo-EM images of P5-1C8 IgG in complex with NP-WT S (g) or NP-JN.1 S (h), obtained at the same IgG concentration (1.4 μM), consistent with the simulation conditions. Scale bar: 100 nm. j) Schematic diagram illustrating the binding process and mechanism between P5-1C8 IgG and either nanoparticle-presented WT or JN.1spikes.

Drawing on both MD simulations and experimental data, we generated a mechanistic diagram summarizing the binding and aggregation processes, highlighting the contrasting behaviors of WT and JN.1 spikes upon interaction with P5-1C8 IgG (Figure 4i). For WT spike, high-avidity interactions yield stable antibody-spike complexes that remain anchored on the nanoparticle surface, leading to saturable binding and the formation of stable, monodisperse, single-nanoparticle assemblies, as consistently observed by ns-EM, cryo-EM and MD simulations. In contrast, NP-JN.1 S undergoes a two-step process: in step 1, rapid on-rates drive immediate binding, with multivalent engagement bridging adjacent nanoparticles and nucleating small clusters; in step 2, cluster growth continues, steric crowding displaces a fraction of initially bound IgG, and aggregates expand with minimal IgG remaining engaged.

This accelerated, cooperative aggregation mechanism compensates for the reduced monovalent affinity, thereby sustaining potent neutralization despite weakened intrinsic binding.

## 3. Discussion

Binding metrics of Fab or full-length IgG to monomeric RBD generally correlate with neutralization IC_50_ values,^[9]^ but such measurements fail to capture the dynamic conformations states of the native spike trimer^[12]^ and the contribution of full-length IgG bivalency.^[32]^ Notably, certain antibodies deviate from this correlation, and the mechanisms underlying these outliers remained largely unexplored.^[9b, 9d, 20d]^ Herein, antibody P5-1C8 represents one such exception, highlighting an alternative model of neutralization. All other characterized JN.1-targeting antibodies show neutralization only when possessing high affinity.^[33]^ To dissect its mechanism, we integrated X-ray crystallography, SPR, negative-stain and cryo-EM, and MD simulations to systematically characterize P5-1C8 IgG binding across spike variants. In both free-solution and nanoparticle-displayed systems, P5-1C8 exhibits sub-picomolar avidity and potent neutralization against WT spike. MD simulations further revealed rapid and saturable binding, leading to the formation of stable, monodisperse, single-nanoparticle assemblies. In contrast, while P5-1C8 binds the JN.1 spike with much lower avidity, it maintains potent neutralizing activity through IgG-driven aggregation. Simulations demonstrated that small clusters gradually expand into large-scale aggregates over time. Together, these insights provide a mechanistic framework for “low-avidity, high-potency” antibodies and underscore the importance of evaluating multivalent aggregation or cross-spike bridging during antibody discovery.

Accurately assessing the relationship between affinity, avidity, and neutralization requires careful consideration of experimental design. For example, in SPR assays, different sensor chips, such as CM5, Streptavidin (SA), and NTA, employ distinct immobilization chemistries that can alter protein density, orientation, and conformation, all of which could influence multivalent binding profiles.^[34]^ In this work, we capturing IgG via Protein A to ensure a consistent Fc-down orientation, preserving unobstructed access to the Fab regions for antigen binding,^[34a, 35]^ thereby enabling reliable assessment of multivalent interactions with nanoparticle-displayed spikes. Alternative scaffolds such as DNA origami also hold promise for presenting multiple spikes with precisely programmed geometry. Thus, selecting an appropriate immobilization strategy is essential for accurately evaluating antigen-antibody multivalency, particularly for antibodies like P5-1C8, whose potency depends on multivalent stabilization rather than high-affinity monovalent binding.

Structural analyses further reveal that P5-1C8 neutralizes SARS-CoV-2 through a distinct mechanism of multivalent aggregation. A similar case is observed for the SARS-CoV-2 antibody P5-1H1, which shows undetectable binding yet high neutralization.^[20d]^ P5-1H1 targets highly overlapping epitopes with a comparable approach angle (Fig S17a, Supporting Information),^[19a]^ and displays a similar time-dependent aggregation pattern (Figure S17b, Supporting Information). More cases of aggregation-driven neutralization by antiviral antibodies has been reported—for example, influenza antibody HC19,^[4b]^ Chikungunya virus antibody CHK-124,^[4a]^ and SARS-CoV-2 antibody W328-6H2,^[16a]^—but those cases involve inherently high affinity and were visualized only as endpoint states by EM. EM captures static snapshots, and despite varying our incubation times down to 5 minutes, we failed to observe intermediate assemblies that could illuminate the 3D binding mode. To bridge that gap, we applied molecular dynamics simulations informed by our own experimental K_D_, structural data, and concentration. In contrast to high-affinity antibodies, whose aggregation is driven by direct antigen-antibody-antigen crosslinking, low-affinity antibodies operate via a distinct mechanism: rapid on-rates enable transient inter-spike bridging, and once initial clusters are nucleated, their growth proceeds largely independently of continued IgG engagement, potentially driven by weak spike-spike interactions that are reinforced when multivalently displayed spikes are brought into close proximity. To our knowledge, this study is the first to dissect the dynamics of low-affinity antibody-mediated aggregation, and we hope it sparks more systematic theoretical and experimental evaluation of multivalent synergy in weakly binding antibodies, enriching both mechanistic understanding and future therapeutic discovery.

## Supporting information

Validation Report

Validation Report

Validation Report

Validation Report

Validation Report

Validation Report

Validation Report

Validation Report

Validation Report

## Acknowledgements

This work was supported by National Key R&D Program of China (2022YFA1206400); Beijing Natural Science Foundation (F251001); the National Natural Science Foundation of China (22277017); Strategic Priority Research Program of Chinese Academy of Sciences (Grant No. XDB0770000). N.L., C.C., Z.L., and Y.Z. are supported by CAMS Innovation Fund for Medical Sciences (CIFMS 2019-I2M-5-018). X.D., L.Y. are supported by the National Science Foundation of China (Grant Nos. 22025302).

## Author Contributions

N.L., P.C., and X.D. contributed equally. Y.Y., Y.Z., L.Z., Q.Z., L.Y., and Z.L. conceived and designed the study. N.L., P.C., and X.D. performed most of the experiments, with assistance from H.X., Z.L., Z.S., J.L., and F.G. N.L. performed biophysical and structural assays, and solved and analyzed the ns-EM and cryo-EM structures of the antibody-spike complex. N.L. solved the crystal structure of WT RBD-P5-1C8 Fab complex with assistance from Z.L., Z.S., F.G., S.F., and Y.Y. Q.Z., P.C, and J.L. performed functional assays including flow cytometry and pseudovirus neutralization assays. N.L. and P.C. concentrated and purified the SARS-CoV-2 JN.1 pseudoviruses. X.D. and L.Y. performed the molecular dynamic simulations. Y.C., J.L., Y.H., G.D., Y.J., L.C., X.N., H.Z., K.Z., Y.D., D.L., X.W., D.H., X.P., and C.C. provided additional technical or intellectual assistance. N.L., P.C., X.D., Z.L., L.Y., Q.Z., L.Z., Y.Z., and Y.Y. had full access to data in the study, generated figures and tables, and took responsibility for the integrity and accuracy of the data presentation. Y.Y. and N.L. wrote the manuscript. All authors reviewed and approved the final version of the manuscript.

## Data Availability Statement

The ns-EM and Cryo-EM density maps generated in this study have been deposited in the Electron Microscopy Data Bank (EMDB) under the following accession codes: EMD-65807 (Immune complex of P5-1C8 Fab binding the RBD of SARS-CoV-2 WT 6p spike protein), EMD-65808 (Immune complex of P5-1C8 IgG binding the RBD of SARS-CoV-2 WT 6p spike protein), EMD-65805 (Immune complex of P5-1C8 Fab binding the RBD of Omicron BA.1 6p spike protein, one Fab bound), EMD-65804 (Immune complex of P5-1C8 Fab binding the RBD of Omicron BA.1 6p spike protein, two Fabs bound), EMD-65806 (Immune complex of P5-1C8 IgG binding the RBD of Omicron BA.1 6p spike protein), EMD-65803 (Immune complex of P5-1C8 Fab binding the RBD of Omicron JN.1 6p spike protein, no Fab bound), EMD-65801 (Cryo-EM structure of SARS-CoV-2 WT 6p spike protein in complex with 1.5 P5-1C8 IgG), EMD-65802 (Cryo-EM structure of SARS-CoV-2 WT 6p spike protein in complex with 1 P5-1C8 IgG).The atomic coordinates for the X-ray crystallography structure are available in the Protein Data Bank under accession code 9K6J (Crystal structure of the complex between SARS-CoV-2 WT RBD and P5-1C8 Fab). SDS-PAGE data, representative micrographs, detailed information on atomic models and EM densities, and other supporting data are provided in the Supplementary Information. All data are available from the corresponding author upon request.

