## supporting information for "IgG-Bridging–Seeded Synergistic Aggregation of SARS-CoV-2 Spikes Underlies Potent Neutralization by A Low-Affinity Antibody"

Experimental Section

*Cell lines*: HEK293T cells (ATCC, CRL-3216; RRID: CVCL_0063) were cultured at 37 ºC with 5% CO_2_ in Dulbecco’s minimal essential medium (DMEM) supplemented with 10% (v/v) heat-inactivated fetal bovine serum (FBS) and 100 U/mL penicillin-streptomycin. FreeStyle 293F cells (Thermo Fisher Scientific, R79007; RRID: CVCL_D603) were maintained at 37 ºC in 5% CO_2_ in SMM 293-TII expression medium (Sino Biological, M293TII). The cell lines were obtained from commercial suppliers and handled under standard aseptic conditions. Throughout the study we observed expected morphology and consistent, reproducible protein expression across independent replicates and batches, with no signs of contamination.

*Protein expression and purification*: Codon-optimized genes encoding trimeric spike proteins of SARS-CoV-2 WT, BA.1, BA.5 and JN.1 were synthesized by Tsingke Co., Ltd. and cloned into the SARS-CoV-2 S HexaPro vector (Addgene: 154754). Each construct encoded residues 1-1208 of spike protein, incorporating six proline substitutions (F817P, A892P, A899P, A942P, K986P, V987P), a “GSAS” substitution at the furin cleavage site (residues 682-685) to enhance stability, and a C-terminal T4 fibritin (folden) trimerization motif, an HRV3C protease cleavage site, 8×HisTag and Twin-Strep Tag. Proteins were expressed in FreeStyle 293F cells and harvested 96-hour post-transfection. Supernatants were clarified and purified by Strep-Tactin Sepharose (YEASEN: #20495ES25), followed by size-exclusion chromatography using a Superose 6 Increase 10/300 GL column (Cytiva: #29091596) in TBS buffer. Purity and molecular weight were assessed by SDS-polyacrylamide gel electrophoresis (SDS-PAGE).

For the receptor-binding domain (RBD) of the SARS-CoV-2 prototype (residues Arg319-Lys529), expression was similarly carried out in FreeStyle 293F cells. The RBD construct included a C-terminal 8×His tag and was purified using Ni-NTA affinity chromatography followed by gel filtration in PBS. Protein purity was confirmed by SDS-PAGE. Detailed characterization of all protein constructs is provided in Table S2 (Supporting Information).

*Antibody and Fab fragment production*: Antibody P5-1C8 is a potent neutralizing mAb initially isolated from COVID-19 convalescents following ancestral SARS-CoV-2 infection.^[1]^ The antibody is covered by patent protection (Patent NO. 2023100460521). The variable regions of its heavy and light chains were cloned into expression vectors containing the human IgG1 constant regions, as previously described. Briefly, plasmids encoding the heavy and light chains were co-transfected into HEK293 F cells using polyethylenimine (PEI, Yeasen). After 96 hours, culture supernatants were harvested, and antibodies were purified using AmMag^TM^ Protein A Magnetic beads (GenScript). Bound antibodies were eluted with glycine buffer (pH 2.0) and further purified by size-exclusion chromatography on a Superdex 200 Increase 10/300 GL column (Cytiva: #28990944).

Fab fragments were generated by papain digestion (Lablead) using an IgG-to-papain ratio of 25:1 (w/w) in digestion buffer (100 mM Tris, 2 mM EDTA, 10 mM L-cysteine) at 37 °C for approximately 5 hours. Fc fragments were removed using Protein A Magnetic beads. The concentrations of IgG and Fab were determined by a Nanodrop 600Plus Spectrophotometer (JIAPENG, Shanghai), and final purification was performed by gel filtration using the same Superdex 200 column. Detailed characterization of the IgG and Fab is summarized in Table S2 (Supporting Information).

*Production of pseudoviruses and neutralization assay*: Pseudoviruses were generated using the wild-type SARS-CoV-2 (GenBank: MN908947.3) and several variants. The Beta variant (Pango lineage B.1.351, GISAID: EPI_ISL_700450) carried 10 spike mutations: L18F, D80A, D215G, 242 244del, S305T, K417N, E484K, N501Y, D614G, and A701V. The Delta variant (Pango lineage B.1.617.2, GISAID: EPI_ISL_1534938) included mutations T19R, G142D, 156-157del, R158G, A222V, L452R, T478K, D614G, P681R, D950N. The BA.1 variant (Pango lineage BA.1, GISAID: EPI_ISL_6752027) contained 32 spike mutations, including A67V, Δ69-70, T95I, G142D/Δ143-145, Δ211/L212I, ins214EPE, G339D, S371L, S373P, S375F, K417N, N440K, G446S, S477N, T478K, E484A, Q493R, G496S, Q498R, N501Y, Y505H, T547K, D614G, H655Y, N679K, P681H, N764K, D796Y, N856K, Q954H, N969K, and L981F. The BA.2.75 spike construct was based on BA.2 with additional mutations W152R, F157L, I210V, G257S, D339H, G446S, N460K, and Q493R (reverted).The BA.5 variants (Pango lineage BA.5, GISAID: EPI_ISL_12559461) was constructed with 30 mutations in the spike such as T19I, 24-26del, A27S, G142D, V213G, G339D, S371F, S373P, S375F, T376A, D405N, R408S, K417N, N440K, L452Q, S477N, T478K, E484A, F486V, Q498R, N501Y, Y505H, D614G, H655Y, N679K, P681H, N764K, D796Y, N969K and Q954H. BF.7 (Pango lineage BF.7, GISAID: EPI_ISL_15429967), BQ.1 (Pango lineage BQ.1, GISAID: EPI_ISL_15458271), BQ.1.1 (Pango lineage BQ.1.1, GISAID: EPI_ISL_15458263) were derived from the BA.5 background and included additional of R346T, or K444T and N460K, or all of the three mutation sites. The XBB variant (Pango lineage XBB, GISAID: EPI_ISL_15601178) featured 13 mutation sites of V83A, Δ145, H146Q, Q183E, V213E, G339H, R346T, L368I, V445P, G446S, N460K, F486S, F490S, and the removal of Q493R, compared to BA.2 variant. Addition of G252V to XBB resulted in XBB.1 (PANGO lineage XBB.1, GISAID: EPI_ISL_15596825). The Omicron JN.1 variant (Pango lineage JN.1, GISAID: EPI_ISL_19019427) was constructed with 61 mutations in the spike including T19I, R21T, L24del, P25del, P26del, A27S, S50L, H69del, V70del, V127F, G142D, Y144del, F157S, R158G, N211del, L212I, V213G, L216F, H245N, A264D, I332V, G339H, K356T, S371F, S373P, S375F, T376A, R403K, D405N, R408S, K417N, N440K, V445H, G446S, N450D, L452W, L455S, N460K, S477N, T478K, N481K, V483del, E484K, F486P, Q498R, N501Y, Y505H, E554K, A570V, D614G, P621S, H655Y, I670V, N679K, P681R, N764K, D796Y, S939F, Q954H, N969K. P1143L.

Pseudoviruses were produced by co-transfecting HEK-293T cells (ATCC) with human immunodeficiency virus backbones expressing firefly luciferase (pNL4-3-R-E-luciferase) and pcDNA3.1 vectors encoding the SARS-CoV-2 spike protein variants. Viral supernatants were harvested 48-72 hours post-transfection, clarified by centrifugation, and stored at -80 °C until use.

Neutralization assays were performed by incubating serial dilutions of antibodies with pseudoviruses at 37 °C for 1 h. The mixture was then added to 293T-hACE2 cells. After 48 h, cells were lysed and luciferase activity was measured. The percentage of neutralization was calculated relative to virus-only control. Antibody dilutions for P5-1C8 IgG or Fab started at 10 μg/mL and were tested against pseudoviruses bearing the spike proteins of SARS-CoV-2 WT, Beta, Delta, BA.1, BA.2.75, BA.5, BF.7, BQ.1, BQ.1.1, XBB, XBB.1, and JN.1. The half-maximal inhibitory concentration (IC_50_) was calculated using a four-parameter dose inhibition response model in Graphpad Prism 8.0.

*Crystallization and data collection*: Fab fragments were mixed with SARS-CoV-2 WT RBD at a molar ratio of 1:2 and incubated overnight at 4 °C. The resulting complexes were further purified by gel-filtration chromatography and concentrated to 10 mg mL^-1^ in TBS buffer for crystallization. Initial crystallization screening was performed at 18 °C in 96-well plates using the micro-volume sitting-drop vapor diffusion method, with crystallization kits listed in Table S5 (Supporting Information). A Mosquito liquid handling robot (TTP Labtech) was used to mix 200 nL of protein solution with 200 nL of reservoir solution. Crystals were initially observed in conditions containing 0.2 M ammonium phosphate dibasic, 20% (w/v) polyethylene glycol (PEG) 3,350, pH 8.0 (PEG/ION NO.44). Subsequent optimization was conducted using the sitting-drop vapor diffusion method in 24-well plates. In this stage, 1 μL of protein solution was mixed with 1 μL of reservoir solution. High-quality crystals of the RBD-Fab complex were obtained in 0.2 M ammonium phosphate dibasic, 28% (w/v) PEG 3,350, pH 8.0. The workflow for crystal optimization procedure is illustrated in Figure S2 (Supporting Information). X-ray diffraction data were collected at the BL02U1 beamline of the Shanghai Synchrotron Research Facility (SSRF) and auto-processed with the HKL3000 software. Data collection and processing statistics are summarized in Table S7 (Supporting Information).

*Structure determination and refinement*: The structure of the SARS-CoV-2 WT RBD-P5-1C8 Fab complex was solved by molecular replacement using PHENIX.^[2]^ Model building and iterative refinement were performed with COOT^[3]^ and PHENIX,^[2]^ respectively. Final Ramachandran statistics: 96.98% favored, 3.02% allowed, and 0.00% outliers for the final model. Refinement statistics are listed in Table S7 (Supporting Information). All structure figures were generated using UCSF ChimeraX.^[4]^

*Negative-stain electron microscopy*: Grid preparation, imaging, and data processing were carried out following established protocols with minor modifications.^[5]^ Briefly, SARS-CoV-2 spike proteins (WT, BA.1, BA.5 or JN.1) were incubated with P5-1C8 IgG or Fab, and P5-1H1 IgG (for BA.5 spike only), at a 3:1 Fab-to-protomer molar ratio at room temperature for 0.5-4 hours. The final concentration of the complexes was approximately ~0.3 µg/µL. Prior to grid application, samples were diluted to 0.015 µg/mL in 1× TBS buffer. A 3.5 µL aliquot was applied to 200-mesh copper grids (Zhongjingkeyi Films Technology Co., Ltd), blotted with filter paper, and stained with 2% (w/v) uranyl acetate. Micrographs were acquired using a JEOL JEM-2100F high-resolution transmission electron microscope operating at 200 kV, with images collected at 30,000× magnification using SerialEM automated software. Image processing and 2D/3D reconstruction were performed using Relion 4.0.1,^[6]^ and visualization and alignment of the final EM maps were carried out in UCSF Chimera and ChimeraX.^[4]^ Negative-stain EM maps have been deposited in the Electron Microscopy Data Bank (EMDB; www.emdataresource.org), with accession codes provided in Table S3 (Supporting Information).

*Cryo-EM sample preparation and data acquisition*: Purified SARS-CoV-2 WT spike protein was mixed with P5-1C8 IgG at a 3:1 molar ratio of IgG to spike trimer and incubated at room temperature for 2 h. The final concentrations of the complex was adjusted to 1 mg/mL in TBS buffer. Prior to vitrification, Quantifoil R1.2/1.3 Cu 300-mesh grids were glow-discharged for 20 s at medium power using a plasma cleaner. A 4 μL aliquot of the complexes was applied to the grids, using a blot force of -2 and a 3-second blot time at 100% relative humidity and 4 °C, and plunge-frozen in liquid ethane using FEI Vitrobot system. Cryo-EM data were acquired on a FEI Titan Krios transmission electron microscope (Thermo Fisher Scientific) operating at 300 kV, equipped with a Gatan K3 Summit direct electron detector. Automated data collection was performed using SerialEM software at 29,000x magnification, corresponding to a pixel size of 0.97 Å. Images were collected at a defocus range of -1.0 to -1.6 μm. Each movie stack was recorded with a total electron dose of 50 e/Å^2^ fractionated in 32 frames.

*Cryo-EM data processing*: Cryo-EM micrographs were processed using CryoSPARC v4.6.0.^[7]^ Contrast transfer function (CTF) parameters were estimated using the Patch CTF Estimation. Initial particle picking was performed with Blob Picker, followed by reference-free 2D classification. Based on these results, the Template Picker was applied to re-pick particles, which were then subjected to a second round of 2D classification. Particles were extracted using a 480-pixel box size and refined through several iterative rounds of 2D classification. Well-defined particles were used for Ab-Initio Reconstruction and Homogeneous Refinement. The resulting dataset was then imported into Relion for 3D classification to eliminate low-quality and heterogeneous particles. Composite masks targeting RBD-Fab and full Spike-Fabs regions were applied during classification to resolve distinct conformational states, including particles with two RBDs in the “up” position and one “down”, or all three RBDs in the “up” position. Subsequently, particles from selected classes were re-imported into CryoSPARC for Local refinement. Final cryo-EM maps were interpreted by fitting atomic models using UCSF Chimera.^[4]^ The RBD was modeled using PDB entry 7JZM, and the P5-1C8 Fab structure was derived from crystallographic data. All cryo-EM visualizations were rendered in Chimera or ChimeraX. Data collection parameters and the complete processing workflow are summarized in Table S6 (Supporting Information) and illustrated in Figure S9 (Supporting Information), respectively.

*Binding of P5-1C8 antibody to cell surface-expressed spike glycoproteins*: HEK293T cells were transfected with plasmids encoding full-length spike proteins of various SARS-CoV-2 variants and incubated at 37 °C for 36 h. Following incubation, cells were detached using trypsin and seeded into 96-well plates. Between each staining step, cells were washed twice with 200 µL of staining buffer (PBS containing 2% heated-inactivated fetal bovine serum). To assess cross-reactive binding, cells were first incubated with the P5-1C8 antibody at a concentration of 5 μg/mL in 100 μL staining buffer for 30 minutes at 4 °C. The tested spike variants included SARS-CoV-2 WT, Beta, Delta, BA.1, BA.2.75, BA.5, BF.7, BQ.1, BQ.1.1, XBB, XBB.1, and JN.1. After washing, cells were stained with PE-conjugated anti-human IgG (H+L) (Biolegend 410708) in 50 μL staining buffer for 30 minutes at 4 °C. Following additional washes, cells were resuspended and analyzed using a BD LSRFortessa flow cytometer (BD Biosciences, USA). Data were processed with FlowJo v10 (FlowJo, USA). Mock-transfected cells served as negative controls. MFI (Mean Fluorescence Intensity) was calculated and exported by Flowjo.

*Surface plasmon resonance (SPR) analysis of IgG and Fab binding to SARS-CoV-2 spike trimer*: Binding kinetics and affinity of P5-1C8 IgG or Fab fragments to SARS-CoV-2 spike trimers (WT, BA.1, and JN.1) were measured using a Biacore 8K SPR system (Cytiva). His-tagged spike trimers were immobilized on a Ni^2+^-charged NTA sensor chip, followed by injection of P5-1C8 IgG or Fab as analytes. NTA surfaces were regenerated with 350 mM EDTA, recharged with 0.5 mM NiCl_2_, and activated with EDC/NHS. Final immobilization levels were ~2900 RU (WT), ~4900 RU (BA.1), and ~6000 RU (JN.1). Binding assays were conducted at a flow rate of 30 μL/min in PBST buffer. Serial dilutions of P5-1C8 IgG or Fab were injected over the immobilized spike protein for 120 s, followed by a dissociation phase of 900 s (WT) or 360 s (BA.1 and JN.1), using a multi-cycle method. The sensor surface was regenerated between cycles with 10 mM glycine-HCl (pH 2.0) for 60 s at a flow rate of 30 μL/min. Kinetic parameters were determined by fitting the reference-subtracted sensorgrams to a 1:1 (Langmuir) binding model (WT and BA.1) or bivalent analyte model (JN.1) using Biacore Insight Evaluation Software (Cytiva).

*Ns-EM analysis of antibody interacting with spike displayed on nanoparticle*: NP-S nanoparticles used in this study were supplied by our collaborator and the formulation procedure, based on a confidentiality agreement, cannot be disclosed in detail. These NP-S nanoparticles were further incubated with P5-1C8 IgG at a 3:1 IgG-to-Spike molar ratio for 5-30 minutes at room temperature, maintaining a final concentration of ~0.3 µg/µL. Prior to grid preparation, samples were diluted to 0.015 µg/µL in 1× TBS buffer. A 3.5 µL aliquot of the diluted sample was applied to 200-mesh copper grids (Zhongjingkeyi Films Technology Co., Ltd), blotted with filter paper, and stained with 2% (w/v) uranyl acetate. Imaging the nanoparticle states and complexes was performed using a transmission electron microscope operated at 80 kV.

*Cryo-EM analysis of antibody interacting with spike displayed on nanoparticle*: Following a similar procedure to the negative-stain protocol, these NP-S nanoparticles were incubated with P5-1C8 IgG at varying IgG-to-Spike molar ratios for 5 minutes at room temperature, maintaining a final concentration of ~0.3 µg/µL. For cryo-EM grid preparation, Quantifoil R1.2/1.3 Cu 300-mesh grids were glow-discharged for 20 s at medium power using a plasma cleaner. A 4 μL aliquot of the sample was applied to each grid, blotted with a force setting of -2 and a 3-second blot time at 100% relative humidity and 4 °C, and plunge-frozen in liquid ethane using a FEI Vitrobot system. Cryo-EM images were acquired on a Thermo Fisher Talos transmission electron microscope operating at 200 kV, using a nominal magnification of 92,000x, which corresponds to a pixel size of 1.59 Å.

*SPR analysis of IgG binding to soluble and nanoparticle-displayed spike proteins*: SPR assays were conducted using a Biacore 8K system (Cytiva) to evaluate the binding kinetics between P5-1C8 IgG and either soluble spike trimers or nanoparticle-displayed spike (NP-S) constructs. Protein A was covalently immobilized onto the CM5 chip via standard amine coupling in 10 mM sodium acetate buffer (pH 4.0), and P5-1C8 IgG was subsequently captured on the Protein A surface. SPR measurements were carried out at a flow rate of 30 μL/min in PBST buffer (PBS with 0.05% Tween-20). Serial dilutions of either soluble spike or NP-S nanoparticles were injected over the sensor surface for 120 s, followed by a dissociation phase of 900 s. The chip surface was regenerated between cycles using 10 mM glycine-HCl (pH 2.0) for 60 s. Sensorgrams were reference-subtracted and analyzed using the Biacore Insight Evaluation Software (Cytiva), and kinetic parameters were obtained by fitting to a 1:1 (Langmuir) binding or bivalent analyte model.

*Negative-stain EM analysis of antibody-induced virus aggregation*: Concentrated SARS-CoV-2 JN.1 pseudovirus were purified by size-exclusion chromatography on a Superose 6 Increase 10/300 GL column (Cytiva, #29091596) equilibrated in PBS. Purified JN.1 pseudovirus was incubated with P5-1C8 IgG for 1 h at room temperature. Subsequently, A 3.5 µL aliquot was applied to 200-mesh copper grids, adsorbed for 5 min, blotted, and stained with 2% (w/v) uranyl acetate, then air-dried. Grids were examined by transmission electron microscope operated at 80 kV, and multiple, randomly selected fields were imaged to assess antibody-induced aggregation. All procedures were conducted under appropriate biosafety conditions.

*Coarse-grained methods, molecular dynamics, and potential energy settings*: Coarse-grained (CG) molecular dynamics (MD) simulations^[8, 9]^ were employed to examine the binding and aggregation behavior of P5-1C8 IgG interacting with either SARS-CoV-2 WT or JN.1 spike trimers displayed on nanoparticles. The CG methodology simplifies the molecular system by representing groups of atoms as single interaction sites, or “beads”, reducing computational complexity while preserving key structural and dynamic features. In this model, each residue of spike and antibody structures were mapped onto one or more CG beads based on its chemical properties, with hydrophobic, polar, and charged side chains grouped accordingly, and the backbone represented by separate beads. The length unit σ was set to 7.5 nm, the time unit $\text{t}_{\text{0}}$ to 0.1 us, and the energy unit ε to $\text{k}_{\text{B}}\text{T}$ at 300 K, where $\text{k}_{\text{B}}$ is the Boltzmann constant ($\text{k}_{\text{B}}\text{=1.38×}\text{10}^{\text{-23}}\text{ }\text{J }\text{K}^{\text{-1}}$):

$\text{ε=}\text{k}_{\text{B}}\text{T}\text{=1.38×}\text{10}^{\text{-23}}\text{J }\text{K}^{\text{-1}}\text{×300K=4.14×}\text{10}^{\text{-21}}\text{J}$	(1)

The molecular dimensions were defined as follows: the nanoparticle radius ($\text{R}_{\text{NP}}$) was set to 7.5 nm, spike trimer length ($\text{L}_{\text{Spike}}$) was 25 nm, and antibody length ($\text{L}_{\text{IgG}}$) was 15 nm. The molecular masses were assigned as $\text{M}_{\text{Spike}}\text{=540 kg }\text{mol}^{\text{-1}}$ for WT or JN.1 spike protein and $\text{M}_{\text{IgG}}\text{=150 kg }\text{mol}^{\text{-1}}$ for antibody.

CG-MD simulations were performed using the GALAMOST software package^[10]^ within the *NVT* ensemble to maintain a constant temperature of 300 K. Temperature control was achieved through a velocity rescaling thermostat with a coupling constant of 0.2 ps, while pressure was regulated using Nose Hoover thermostat with a coupling constant of 1 ps and a thermostat coupling parameter of 0.5. For simulations involving spike trimers displayed on nanoparticles, a cubic box with a fixed dimensions of 1500×1500×1500 nm^3^ was used to accommodate a large number of nanoparticles. To better simulate experimental conditions, the system included ~0.5 μM spike trimer displayed on nanoparticles and 1.4 μM IgG, with parameters and methodologies ensuring representation of the essential system physics. The equations of motion were integrated using the Velocity-Verlet algorithm with a time step of 4 ns. Each simulation was run for a total duration of 15 μs, a timescale sufficient to capture both short-term binding events and long-term aggregation processes. Trajectories were saved at 1 ns interval for subsequent analysis.

Interactions between CG beads were described using a combination of bonded and non-bonded potentials. Bonded potentials included harmonic terms for bond lengths and angles, as well as dihedral potentials to maintain the overall molecular structure. The bond length potential was typically modeled using a harmonic potential, which can be expressed as:

$\text{U}_{\text{b}\text{ond}}\left( \text{r} \right)\text{=}\frac{\text{1}}{\text{2}}\text{k}_{\text{b}}\left( \text{r}\text{-}\text{r}_{\text{0}} \right)^{\text{2}}$	(2)

where $\text{U}_{\text{b}\text{ond}}\left( \text{r} \right)$ represents the potential energy as a function of bond length, *k*_b_ is the force constant that determines the bond stiffness, and $\text{r}_{\text{0}}$ is the equilibrium bond length. This harmonic potential ensures that the distance between two bonded beads remains close to the equilibrium value $\text{r}_{\text{0}}$, with deviations leading to an increase in potential energy. The bond angle potential is typically modeled using a harmonic potential:

$\text{U}_{\text{angle}}\left( \text{θ} \right)\text{=}\frac{\text{1}}{\text{2}}\text{k}_{\text{θ}}\left( \text{θ}\text{-}\text{θ}_{\text{0}} \right)^{\text{2}}$	(3)

where $\text{U}_{\text{angle}}\left( \text{θ} \right)$ is the potential energy as a function of bond angle $\text{θ}$, $\text{k}_{\text{θ}}$ is the force constant for angle, and $\text{θ}_{\text{0}}$ is the equilibrium bond angle. This potential maintains the preferred angle between three consecutive beads, thereby contributing to the overall conformational stability of the molecule.

Non-bonded potentials describe the interactions between beads that are not directly connected, including van der Waals (VDW) and electrostatic interactions. The Lennard-Jones (LJ) potential is commonly used to model van der Waals interactions:

$\text{U}_{\text{L}\text{J}}\left( \text{r} \right)\text{ = 4ε}\left[ \left( \frac{\text{σ}}{\text{r}} \right)^{\text{1}\text{2}}-\left( \frac{\text{σ}}{\text{r}} \right)^{\text{6}} \right]$	(4)

where $\text{U}_{\text{L}\text{J}}\left( \text{r} \right)$ is the potential energy as a function of the distance *r* between two CG beads, ε is the potential well depth representing the interaction strength, and σ is the distance at which the potential energy is zero, related to the size of the interacting beads. The LJ potential captures both the attractive and repulsive forces between non-bonded beads, providing a realistic description of van der Waals interactions. Electrostatic interactions are typically modeled using the Coulomb potential:

$\text{U}_{\text{C}\text{oulomb}}\left( \text{r} \right)\text{=}\frac{\text{q}_{\text{i}}\text{q}_{\text{j}}}{\text{4}\text{π}\text{ϵ}_{\text{0}}\text{ϵ}_{\text{r}}\text{r}}$	(5)

where $\text{U}_{\text{C}\text{oulomb}}\left( \text{r} \right)$ is the potential energy as a function of the distance *r* between two charged beads, $\epsilon_{0}$is the permittivity of free space, and $\epsilon_{r}$is the relative permittivity (dielectric constant) of the medium. In this study, non-bonded interactions were modeled using Lennard-Jones potential for van der Waals forces and Coulomb potential for electrostatic interactions, parameterized based on the MARTINI force field. Electrostatic interactions were treated using the reaction field method to account for the screening effect of solvent. To clarify, the full simulation parameters, along with their values and sources, are summarized in Table S8 (Supporting Information).

*Analysis of the CG-MD simulations*: All analyses were carried out by means of house-made python scripts employing the Scipy,^[11]^ Pytorch,^[12]^ Scikit-learn,^[13]^ and Fresnel (https://fresnel.readthedocs.io/) package. Snapshots in Figure 4b and Movie S1 and S2 were rendered with Fresnel package using GPU acceleration. For the binding analysis of nanoparticle and antibody, we proceeded as follows: two spikes and antibody were considered in contact (i.e., they belonged to the same assembly) if any of their CG beads lied within a distance of $\text{r}_{\text{cut}}\text{=}\text{ }\text{s}\text{ }\text{=}\text{ }\text{7.5 nm}$. A nanoparticle was considered to be bound if at least one of its grafted spikes contacted an antibody. For aggregation analysis, cluster identification was performed using the DBSCAN algorithm in SciPy. The epsilon parameter, defining the maximum neighbor distance between nanoparticles, was set to 30.0 nm, and the minimum cluster occupancy of 2 was specified to classify even the smallest aggregates while designating isolated points as noise. This density-based approach effectively identifies clusters of arbitrary shapes within simulation trajectories, providing key insights into oligomerization and aggregation processes in simulation.

*Quantification and statistical analysis*: Protein concentrations were determined using a Nanodrop spectrophotometer. SDS-PAGE gels, size-exclusion chromatography profiles, and negative-stain EM micrographs of purified IgG, Fab, spike proteins, and RBD are summarized in Table S2. Neutralization assays were performed in two independent experiments, each with duplicate measurements. Half-maximal inhibitory concentration (IC_50_) were calculated by the equation of four-parameter dose inhibition response using Graphpad Prism 8.0. SPR binding kinetics were conducted in triplicate, and sensorgrams were reference-subtracted and fitted using appropriate kinetic models using Biacore Insight Evaluation 5.0.18.22102. For ns-EM and cryo-EM analyses, the number of micrographs, initial and final particle counts, and class distributions are summarized in Table S3 and Table S6. Data processing and visualization were carried out using CryoSPARC v4.6.0, Relion v4.0.1, Phenix, Coot, UCSF Chimera, UCSF ChimeraX, Biacore Insight Evaluation 5.0.18.22102, Origin 2018, GraphPad Prism v8.0, FlowJo v10, and custom Python scripts employing SciPy, PyTorch, Scikit-learn, and Fresnel libraries. Coarse-grained MD simulations were performed with the GALAMOST package. EM density maps and crystallographic model have been deposited in EMDB and PDB.

Supporting Figures

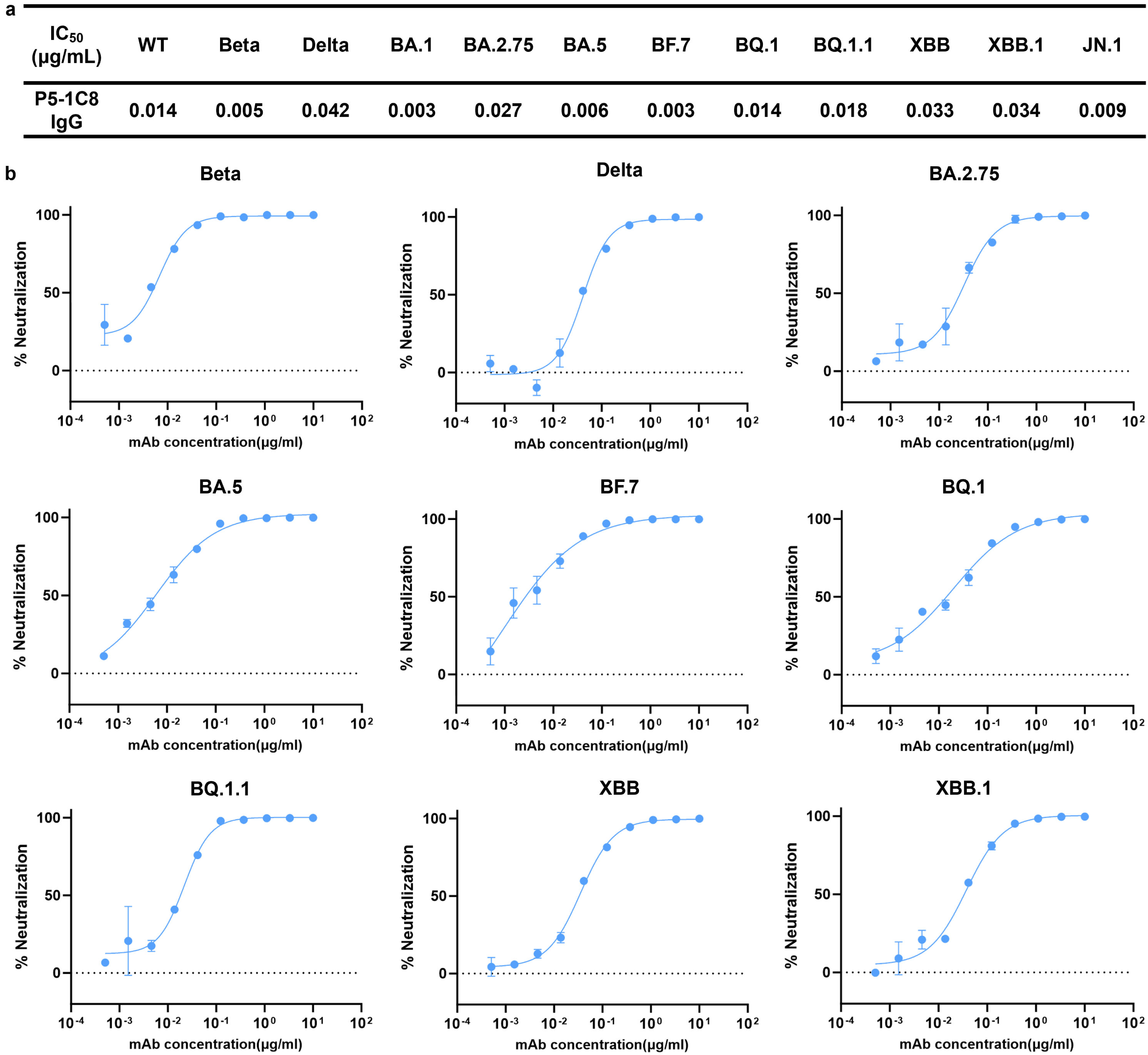
**Figure S1.** Neutralizing capability of P5-1C8 IgG against SARS-CoV-2 variants. a) Neutralizing potency (IC_50_) and neutralization curves (b) of P5-1C8 IgG against a panel of nine coronaviruses, with mean values and standard errors (s.e.m.) indicated. The detection limit, defined by the highest antibody concentration used, was 10 μg/mL. Data was obtained from two independent experiments, each performed with two technical replicates.

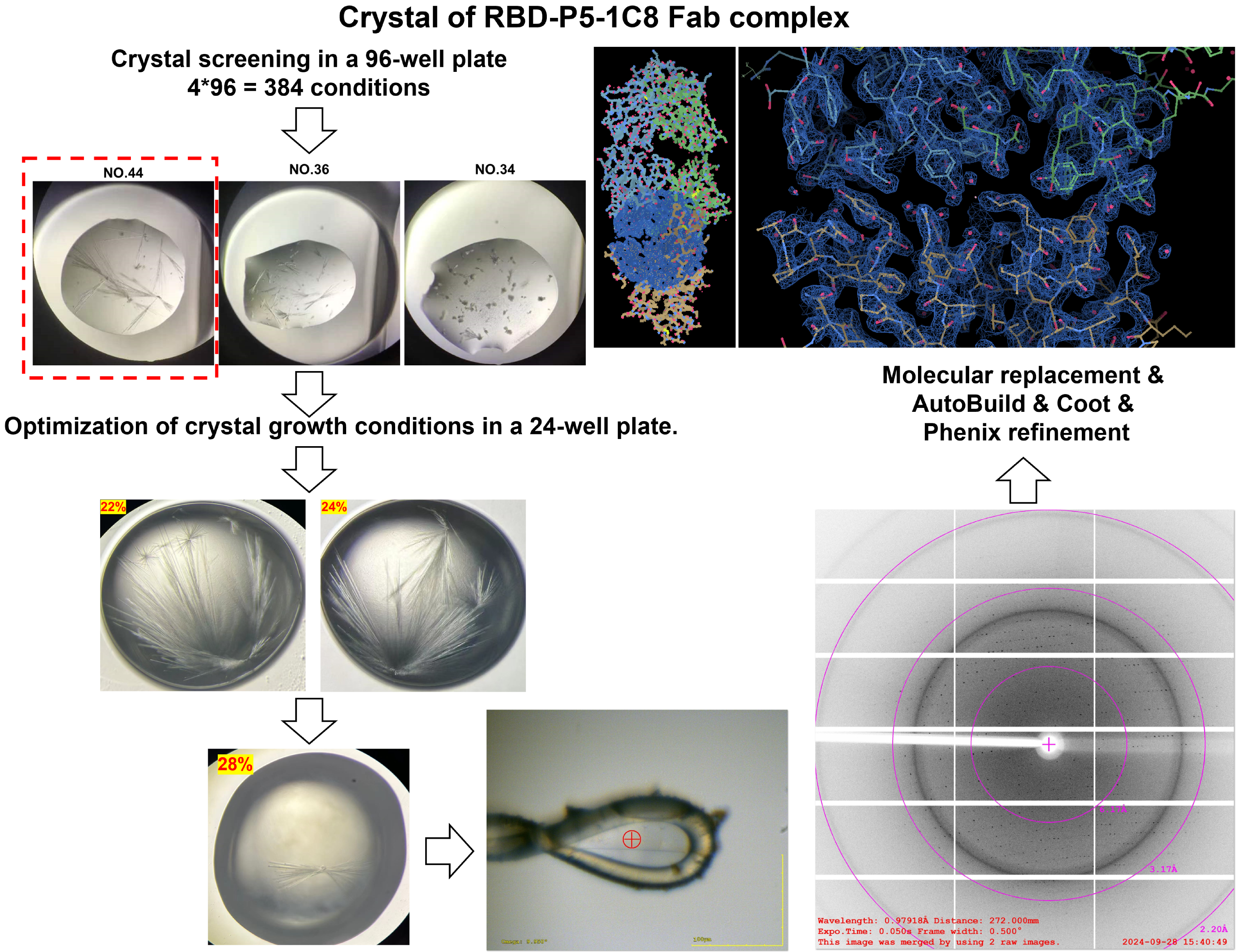
**Figure S2.** Workflow for crystallization optimization, X-ray data collection, and processing of the SARS-CoV-2 wild-type RBD in complex with P5-1C8 Fab.

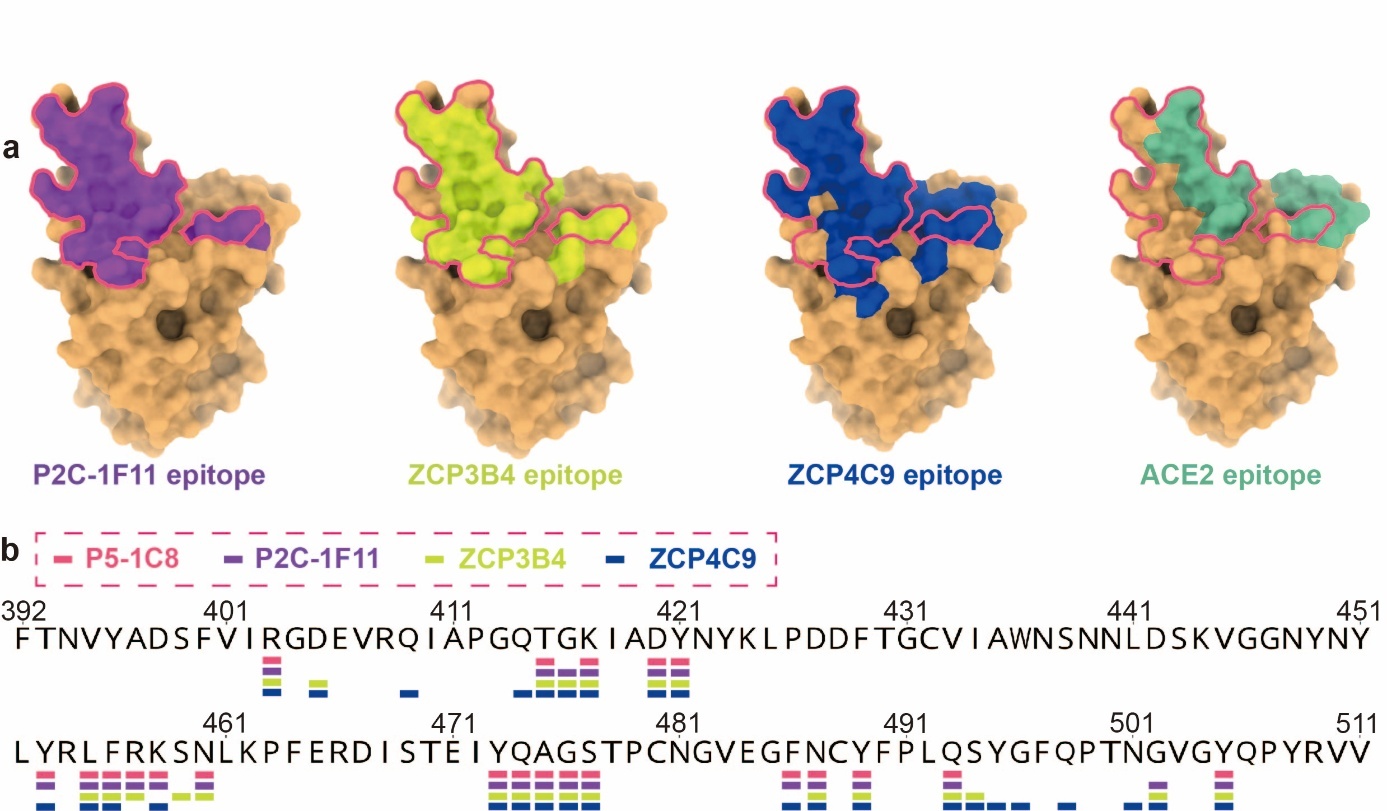
**Figure S3.** Epitope mapping of four class 1 antibodies and ACE2. a) The footprints of three Fabs and ACE2 on SARS-CoV-2 RBD. Dark violet, moderate yellow, blue, and green represent the footprint of P2C-1F11 Fab, ZCP3B4 Fab, ZCP4C9 Fab, and ACE2 respectively. b) RBD sequence from residues 392 to 511, with binding interface residues of the four antibodies highlighted by pink, dark violet, moderate yellow, and blue boxes, respectively.

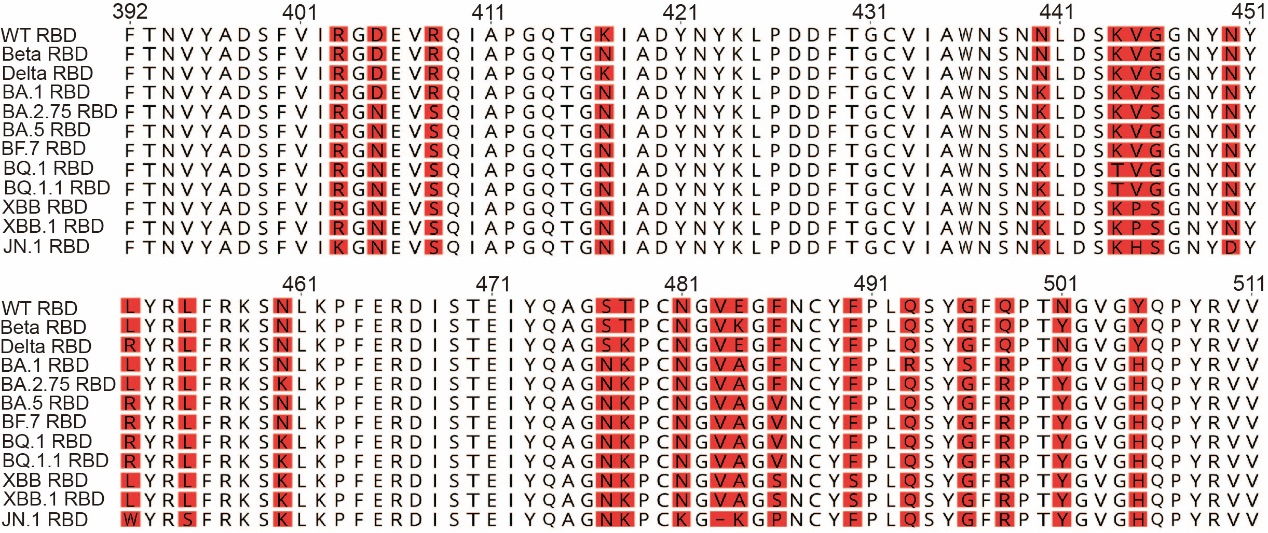
**Figure S4.** Sequence alignment of RBDs from SARS-CoV-2 variants included in the neutralization assays. Amino acids residues differing from the WT RBD are highlighted in red. The RBD sequences correspond exactly to the variant panel used in neutralization assays, including WT, Beta, Delta, BA.1, BA.2.75, BA.5, BF.7, BQ.1, BQ.1.1, XBB, XBB.1, and JN.1.

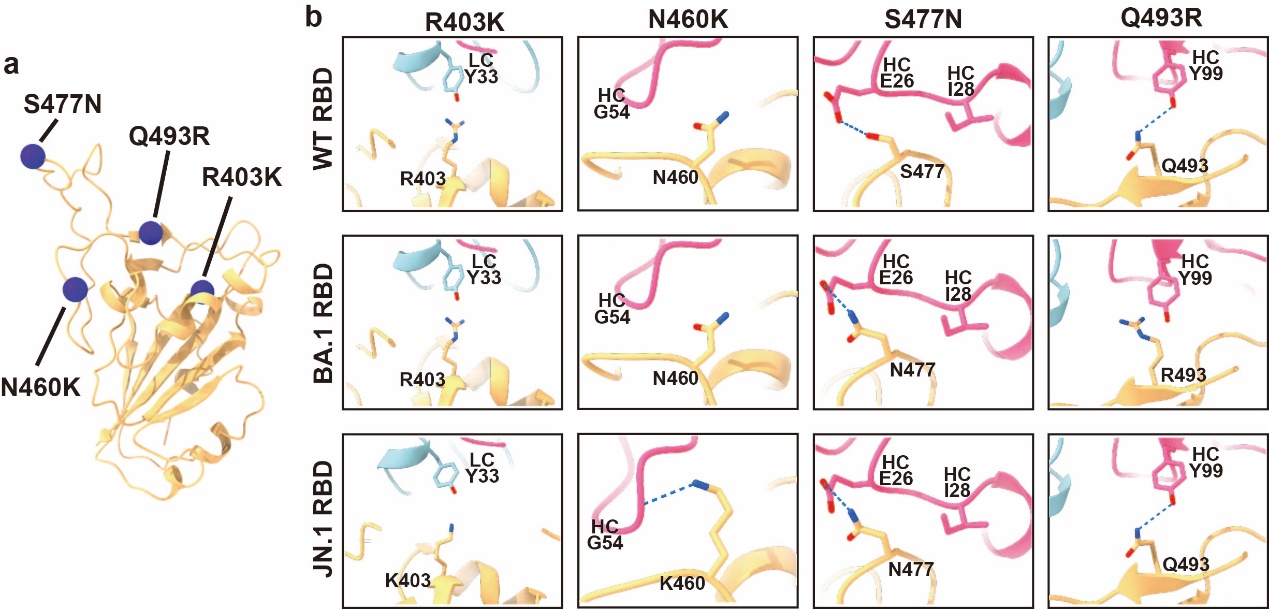
**Figure S5.** The interactions between P5-1C8 Fab and non-conserved residues in the RBD show only minor changes. a) Structure of the SARS-CoV-2 WT RBD, with key P5-1C8 Fab-binding residues highlighted as violet spheres. b) Shown are the structural comparisons of P5-1C8 Fab interactions with WT RBD (upper panel) and Omicron BA.1 RBD (middle panel), and JN.1 RBD (lower panel) at positions R403K, N460K, S477N, and Q493R. Contacting residues are depicted as sticks, and hydrogen bonds or salt bridges are indicated by dashed lines.

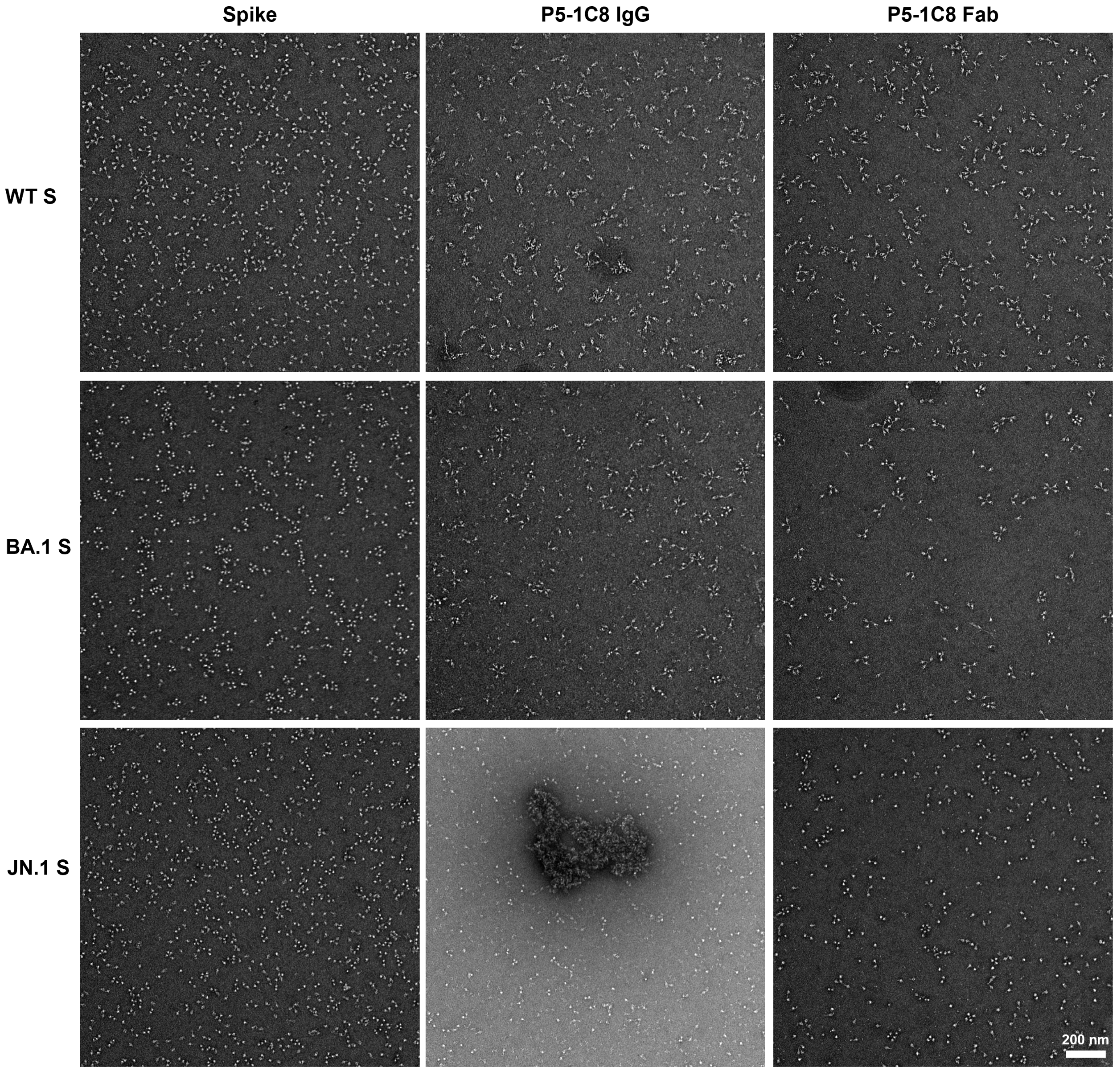
**Figure S6.** Negative-stain micrographs of SARS-CoV-2 spike trimers alone or in complex with P5-1C8 IgG or Fab for WT, BA.1, and JN.1 S. Prior to antibody addition, each soluble spikes were confirmed to be well-dispersed with no detectable aggregation. WT, BA.1, and JN.1 spike trimers were incubated with 3-fold molar excess of P5-1C8 IgG or Fab at RT for 1 h. Scale bar: 200 nm. The scale bar in the last image is applicable to all other images in the same panel.

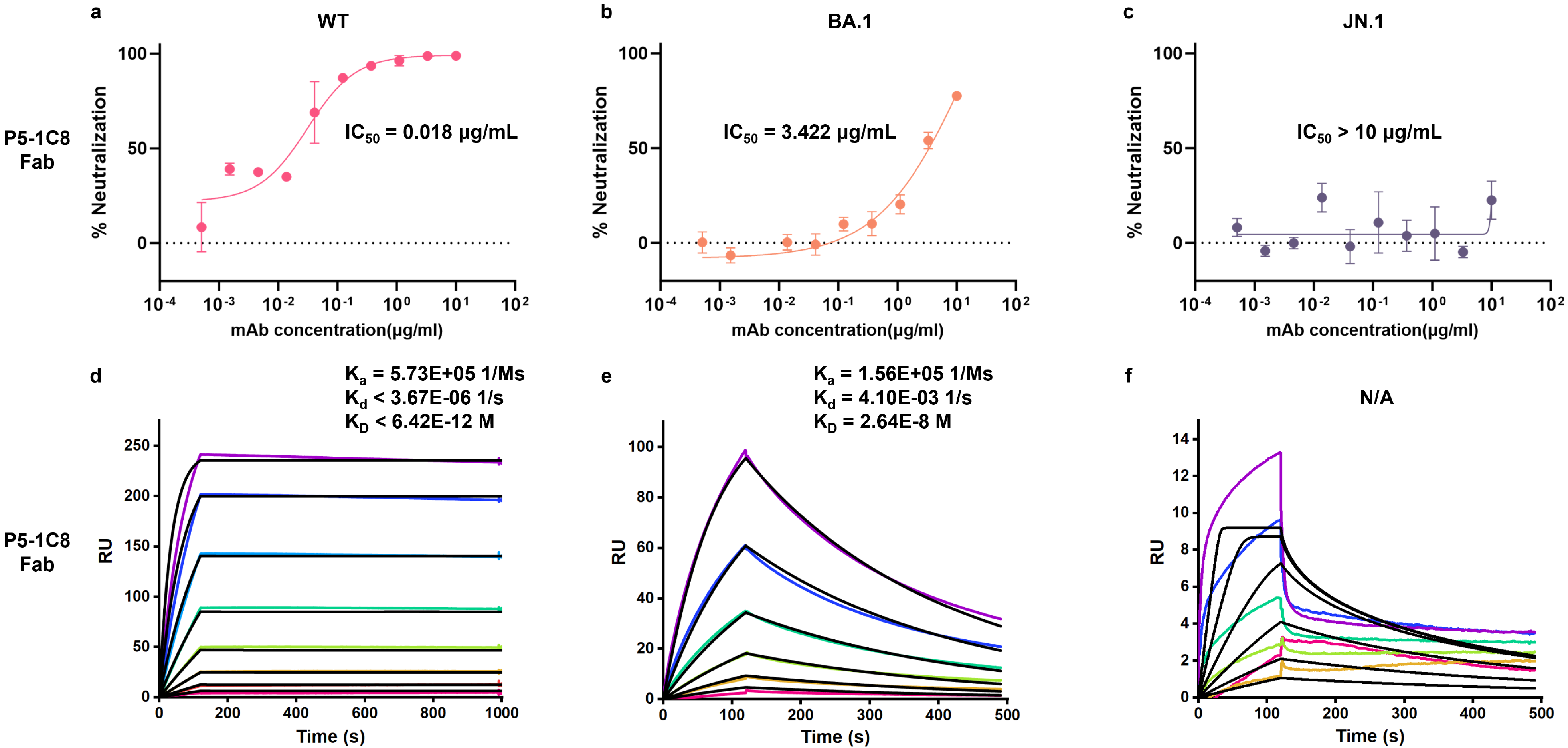
**Figure S7.** Neutralization potency and binding affinity of P5-1C8 Fab. a-c) Neutralization curves of P5-1C8 Fab against SARS-CoV-2 WT (a), BA.1 (b), and JN.1 (c) variants. Data was obtained from two independent experiments, each performed with two technical replicates. d-f) Binding kinetics between P5-1C8 Fab and the spike trimer of SARS-CoV-2 WT (d), BA.1 (e), and JN.1 (f), measured by surface plasmon resonance (SPR). Spike trimers were immobilized on a nitrilotriacetic acid (NTA) sensor chip, and serial dilutions of P5-1C8 Fab were flowed through the system. Colored lines represent experimentally measured sensorgrams. Black lines show the best-fit curves based on experimental data. The calculated association rate (K_a_), dissociation rate (K_d_), and equilibrium dissociation constant (K_D_) for each antibody-spike pair are indicated. All results were confirmed in three independent experiments.

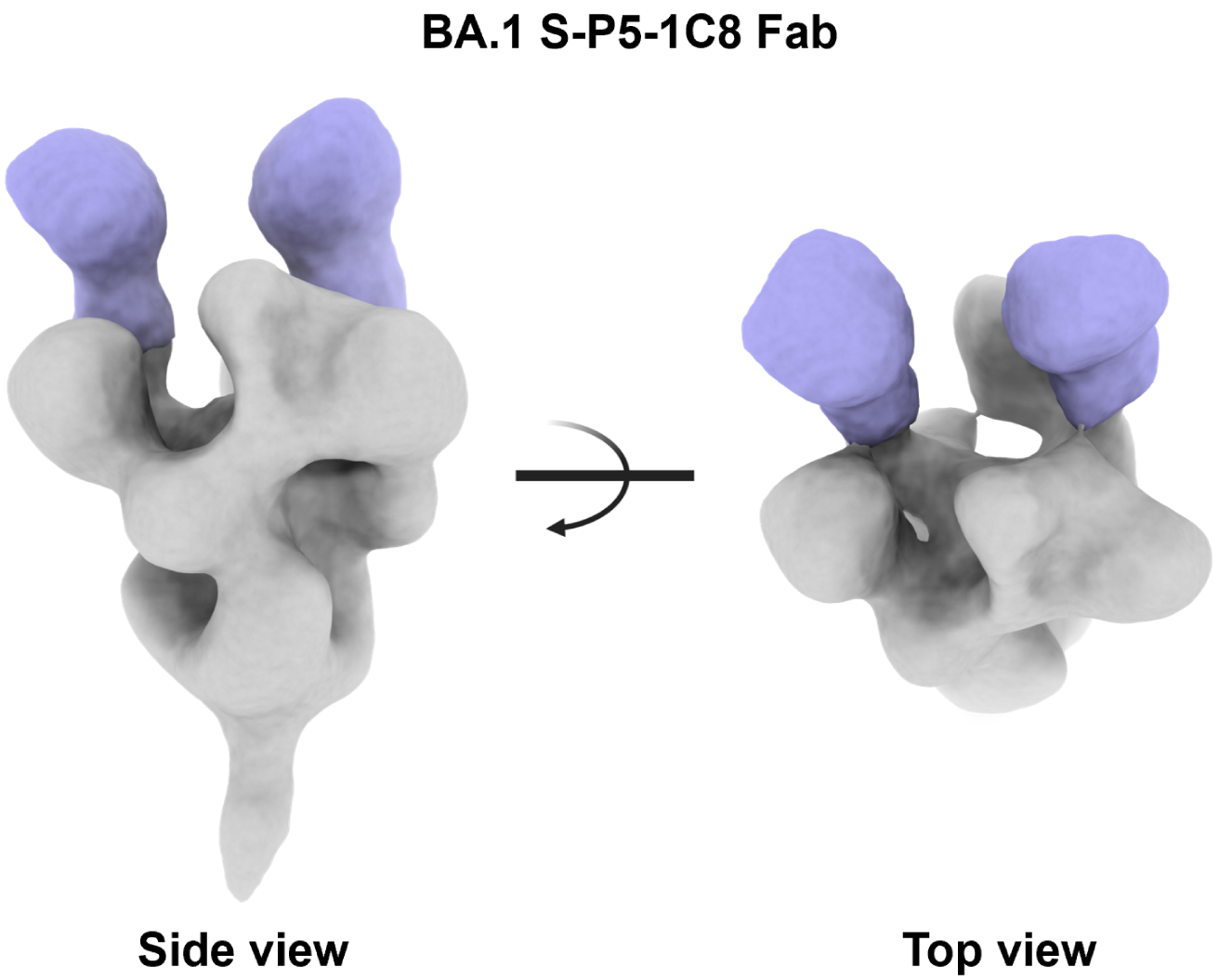
**Figure S8.** Representative 3D reconstructions of Omicron BA.1 spike trimers incubated with P5-1C8 Fab at a 3:1 Fab-to-protomer molar ratio, showing two Fab molecules bound per trimer.

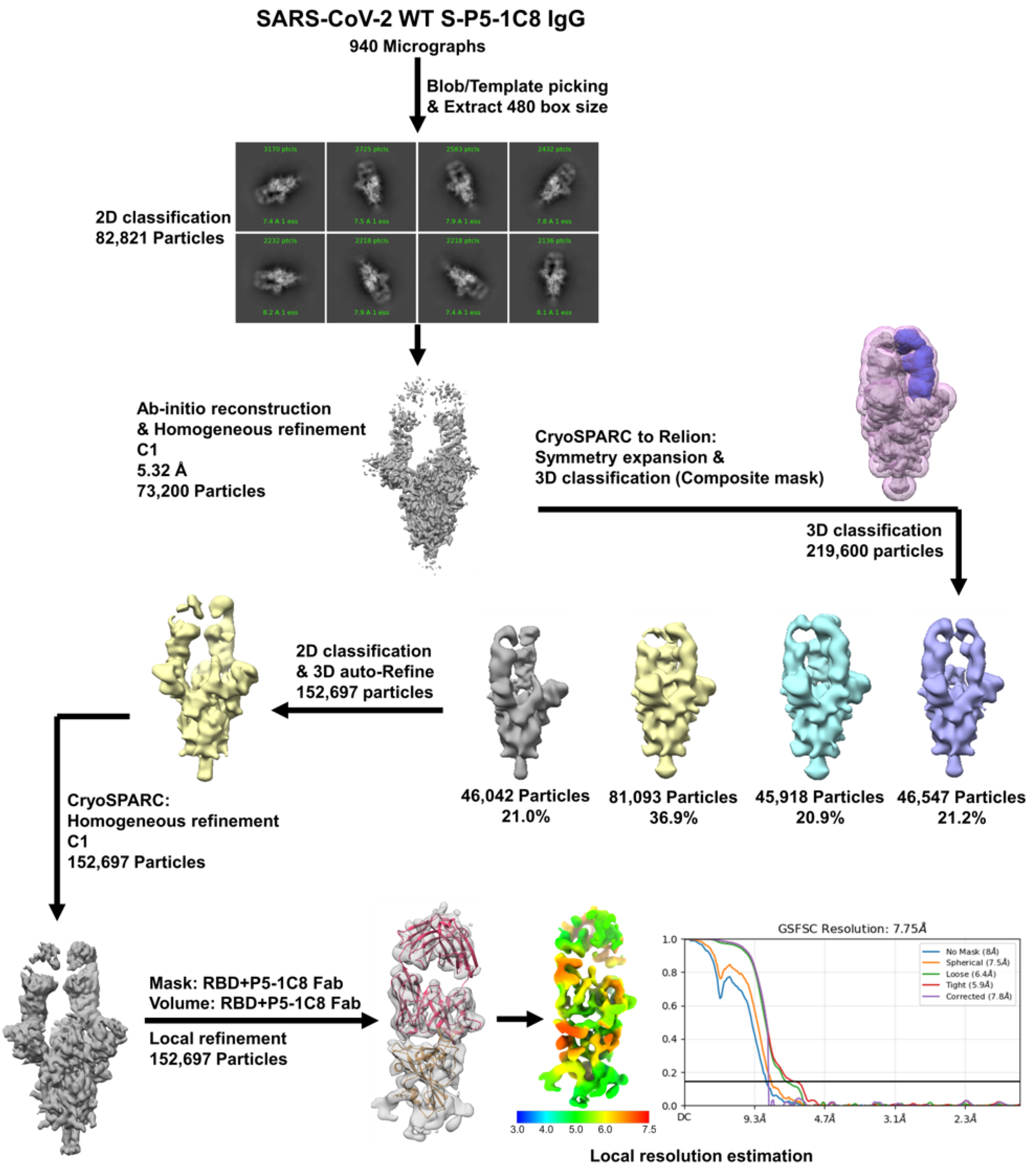
**Figure S9.** Cryo-EM data processing workflow for the SARS-CoV-2 WT spike protein in complex with P5-1C8 IgG.

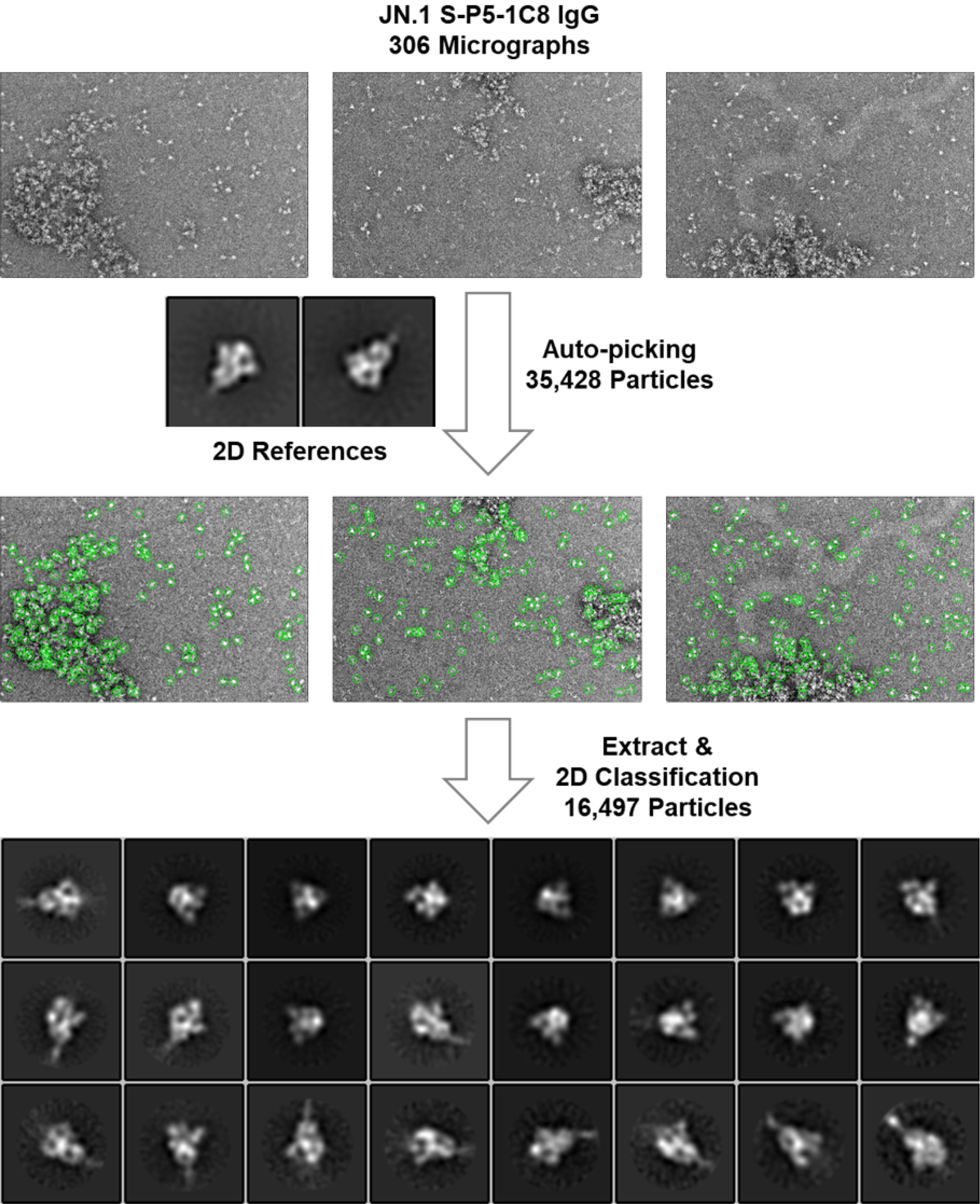
**Figure S10.** Auto-picking and data processing workflow for the JN.1 S-P5-1C8 IgG complex using Relion. Spike only were used as references during auto-picking to select non-aggregated particles. However, 2D class averages showed no detectable IgG density on the spike.

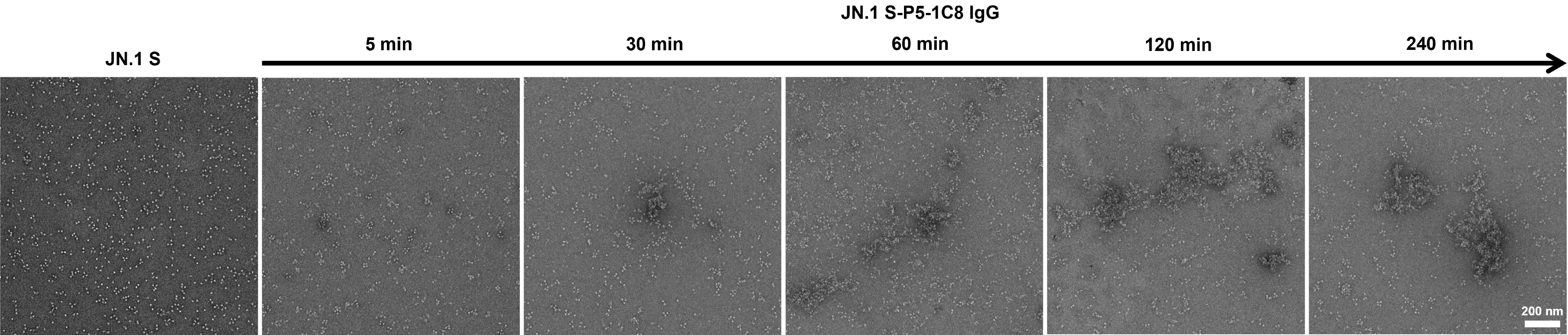
**Figure S11.** Negative-stain EM images of P5-1C8 IgG in complex with JN.1 spike trimers after incubation for 5 to 240 mins. Scale bar: 200 nm. The scale bar in the last image is applicable to all other images in the same panel.

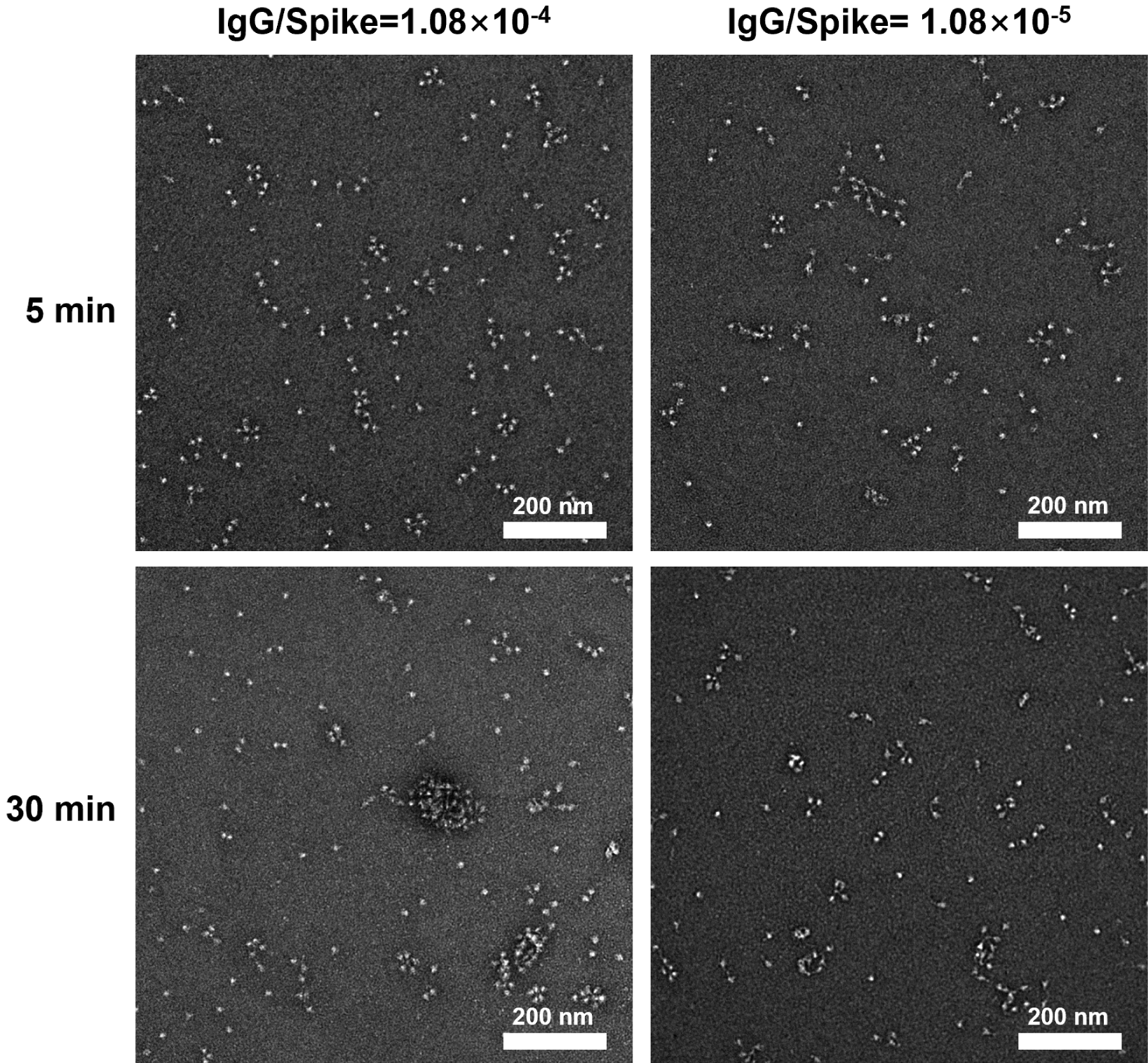
**Figure S12.** Negative-stain EM images of P5-1C8 IgG in complex with JN.1 spike trimers at IgG-to-spike molar ratio ranging from 1.08 × 10^-4^ to 1.08×10^-5^, following incubation for 5 to 30 min. An IgG/Spike ratio of 1.08 × 10^-4^ corresponds to an IgG concentration of 0.06 nM, approximating the IC_50_. Scale bar: 200 nm.

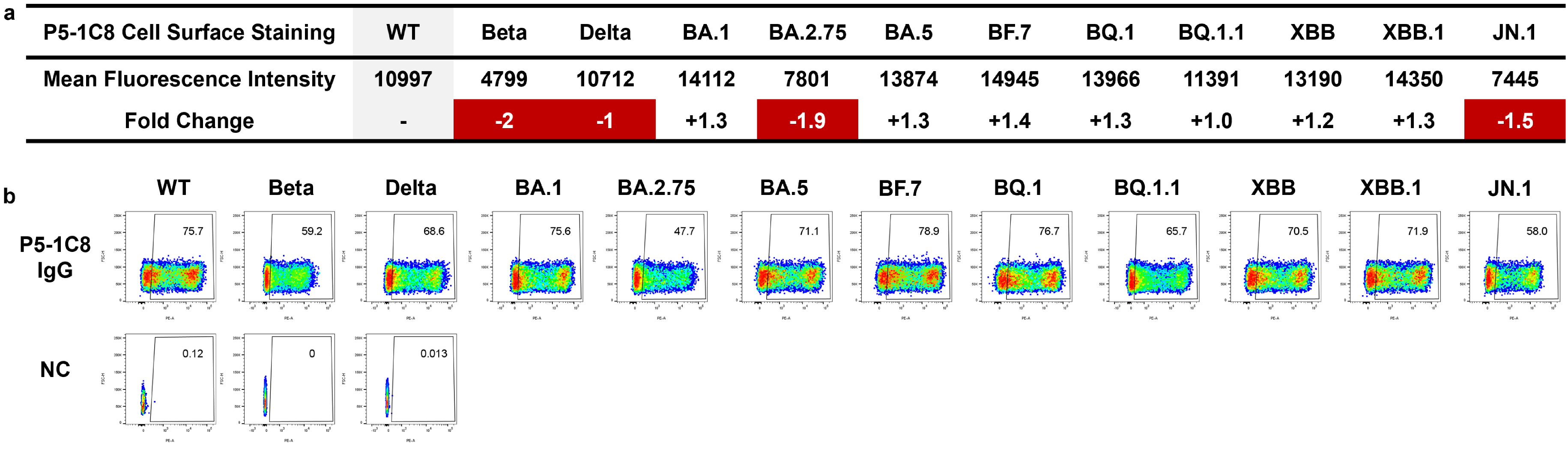
**Figure S13.** Cross-binding ability of P5-1C8 IgG to SARS-CoV-2 variants assessed by cell surface staining. a) For cell surface staining, spike proteins from diverse SARS-CoV-2 variants were expressed on the surface of HEK293T cells, stained with P5-1C8 IgG, and analyzed by flow cytometry. Total fluorescence intensity of positively stained cells was quantified and normalized to represent relative binding ability. Fold change was calculated relative to WT. b) Gating strategies and representative flow cytometry results for P5-1C8 IgG cell surface staining. Numbers in the upper-right corners of each gate indicate the percentage of cells detected by the antibody. Data are representative of two independent experiments.

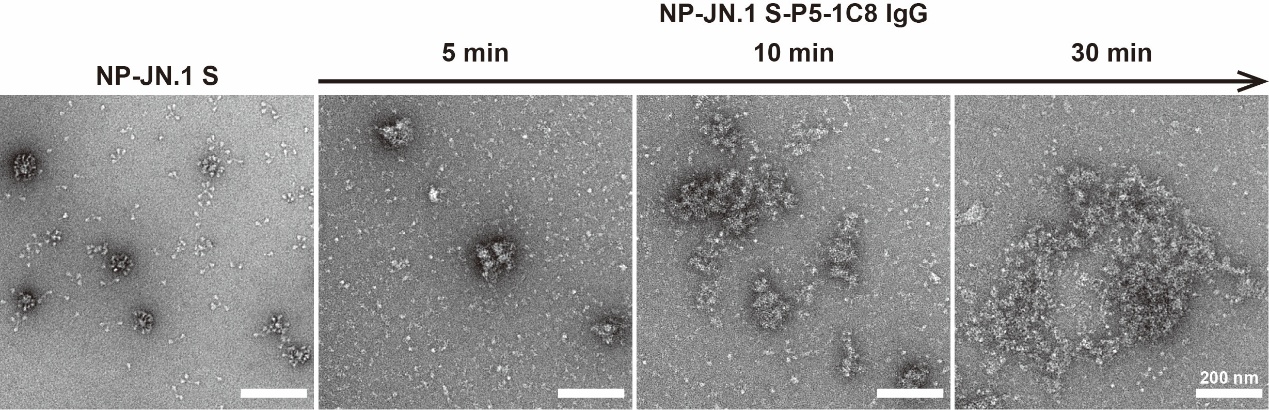
**Figure S14.** Negative-stain EM images of P5-1C8 IgG in complex with NP-JN.1 S nanoparticles after incubation for 5 to 30 min, with IgG concentration of 1.4 μM and ~0.5 μM spike trimer displayed on nanoparticles. Scale bar: 200 nm. The scale bar in the final image is representative of all images in the panel.

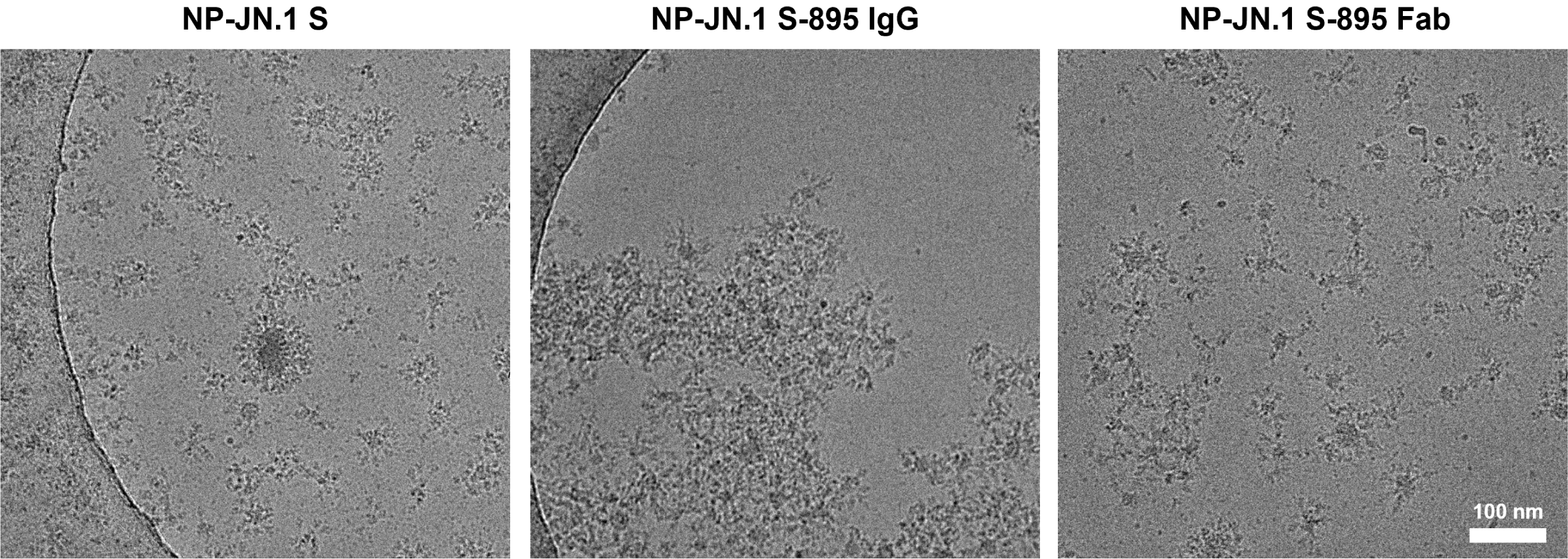
**Figure S15.** Cryo-EM images of P5-1C8 IgG or Fab in complex with NP-JN.1 S nanoparticles at an IgG concentration of 0.2 μM, with ~0.5 μM spike trimer displayed on nanoparticles. Scale bar: 100 nm. The scale bar in the final image represents all images in the panel.

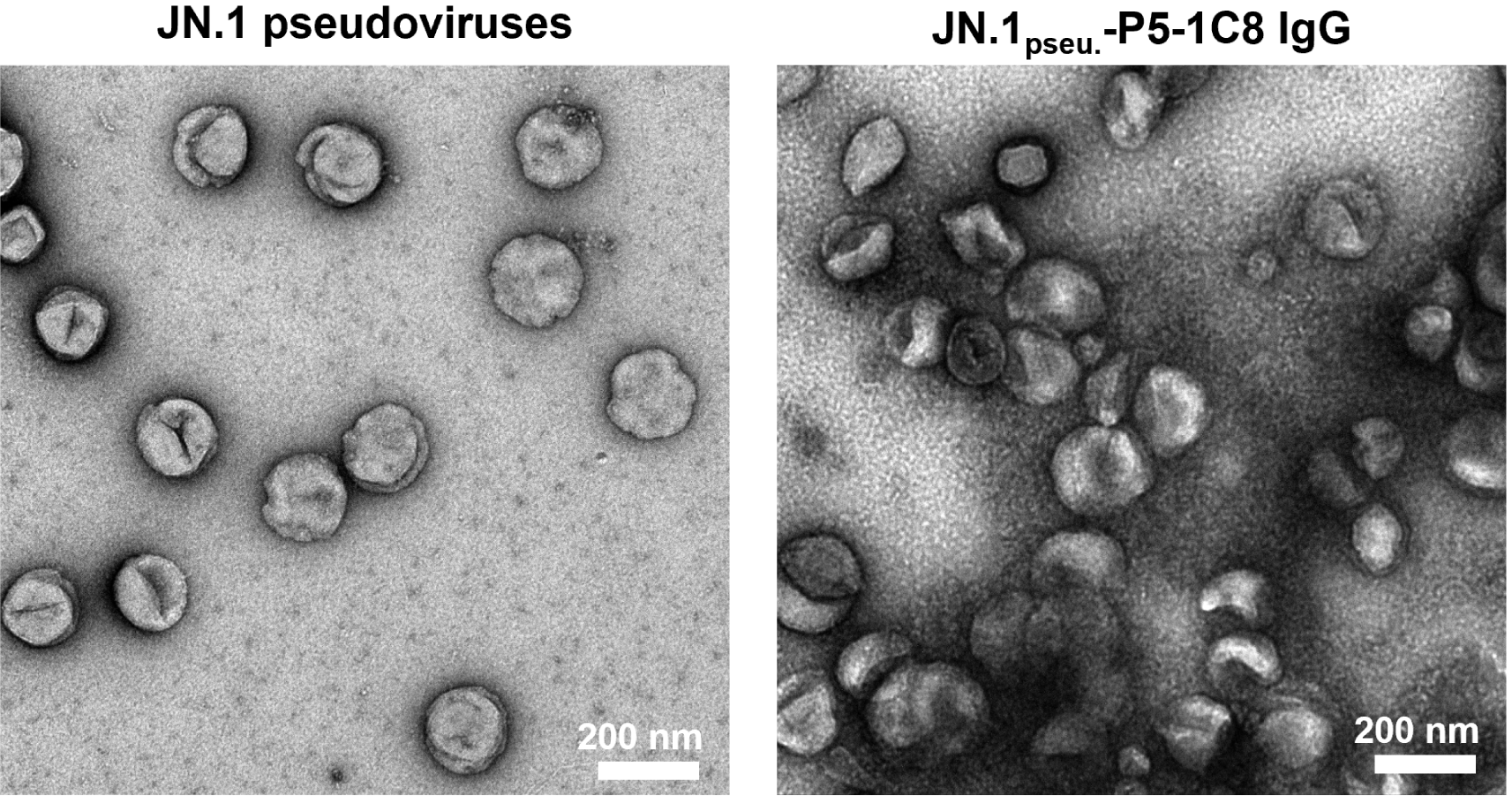
**Figure S16.** Representative negative-stain EM images of JN.1 pseudoviruses and JN.1_pseu._-P5-1C8 IgG complex. Scale bar: 200 nm.

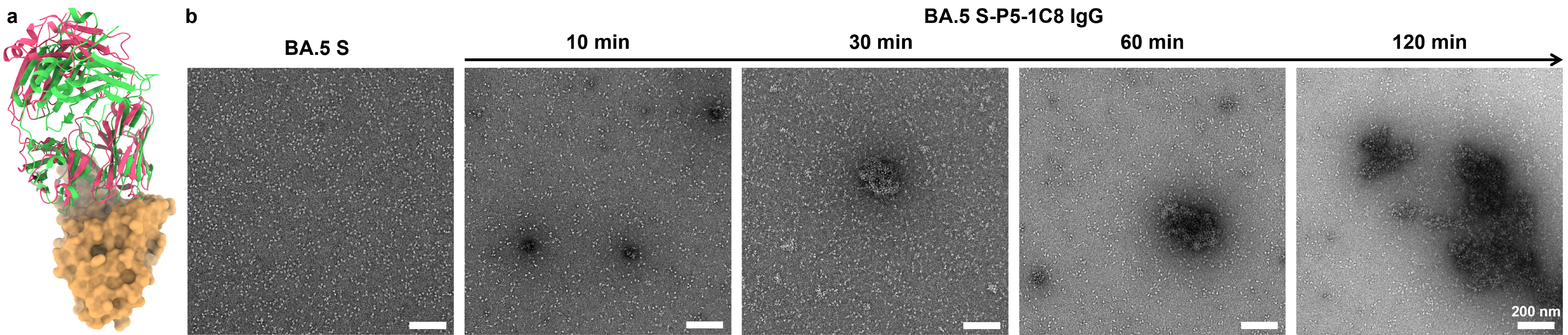
**Figure S17.** Structural comparison and negative-stain EM of P5-1H1 and BA.5 spike trimers interactions. a) Superimposed structures of P5-1H1 Fab (green) and P5-1C8 Fab (pink) bound to the SARS-CoV-2 RBD (soft orange), revealing highly overlapping epitopes and nearly identical approach angles (P5-1H1: PDB 7XS8; P5-1C8: PDB 9K6J). b) Representative negative-stain EM images of P5-1C8 IgG complexed with BA.5 spike trimers after incubation for 10 to 120 mins. Scale bar: 200 nm. The scale bar in the last image applies to all other panels.

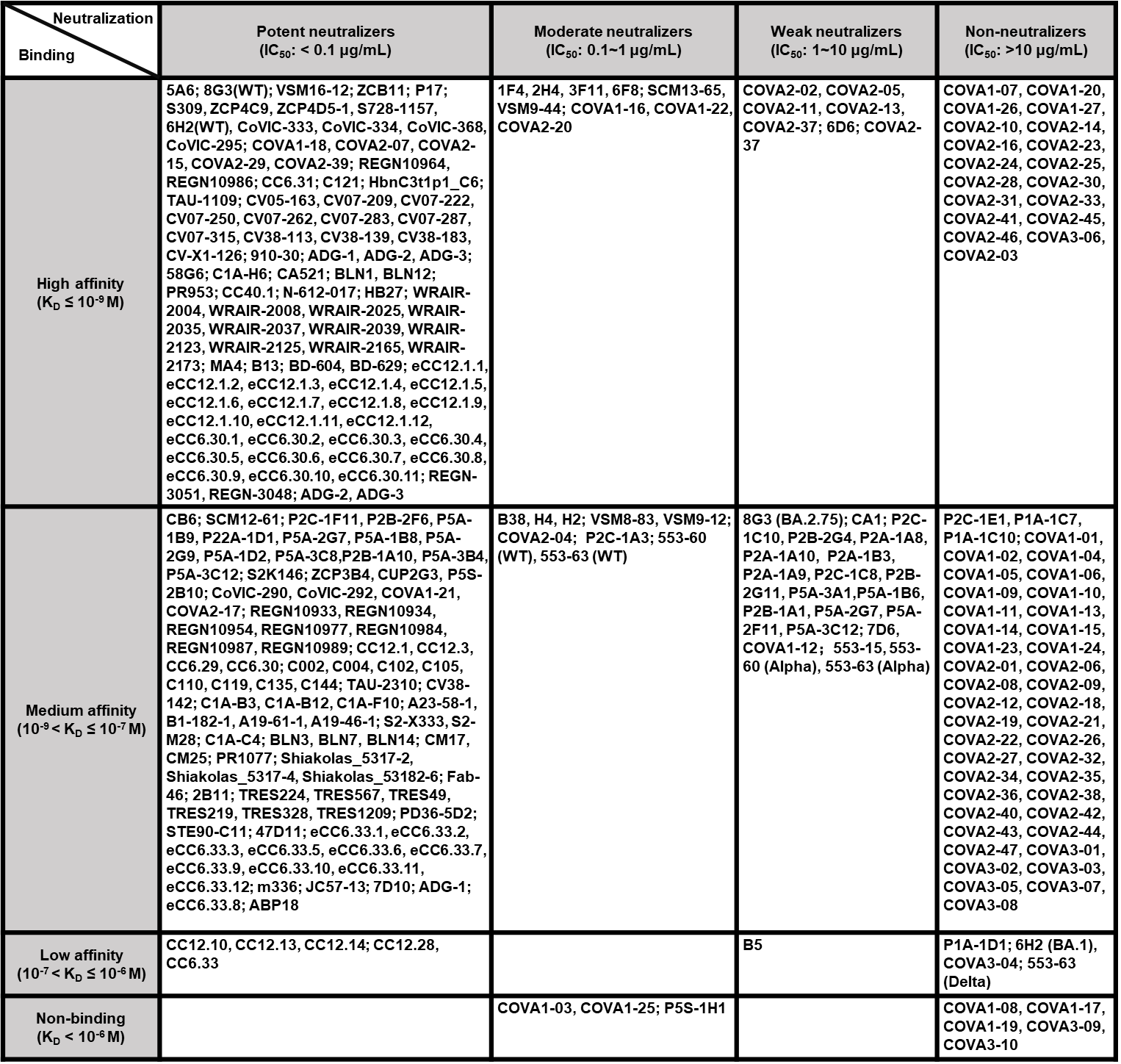
**Table S1.** Correlation of binding affinity and neutralization potency for previously reported SARS‑CoV‑2 antibodies. K_D_ values (measured by SPR or BLI) and IC_50_ values were sourced from the Ab‑CoV database (antibodies reported prior to 2022)^[14]^ and from peer‑reviewed publications in recent years. Antibodies are categorized by affinity^[15]^—high (K_D_ ≤ 10⁻⁹ M), medium (10⁻⁹ < K_D_ ≤ 10⁻⁷ M), low (10⁻⁷ < K_D_ ≤ 10⁻⁶ M), and non‑binder (K_D_ > 10⁻⁶ M or undetectable)—and by neutralization potency^[16]^—potent (IC_50_ < 0.1 μg/mL), moderate (0.1-1 μg/mL), weak (1-10 μg/mL), and non‑neutralizing (IC_50_ > 10 μg/mL).

**Table S2.** SDS-PAGE gels, size-exclusion chromatography profiles, and negative-stain EM micrographs of purified IgG, Fab, spike proteins, and RBD. N.A.: not available.

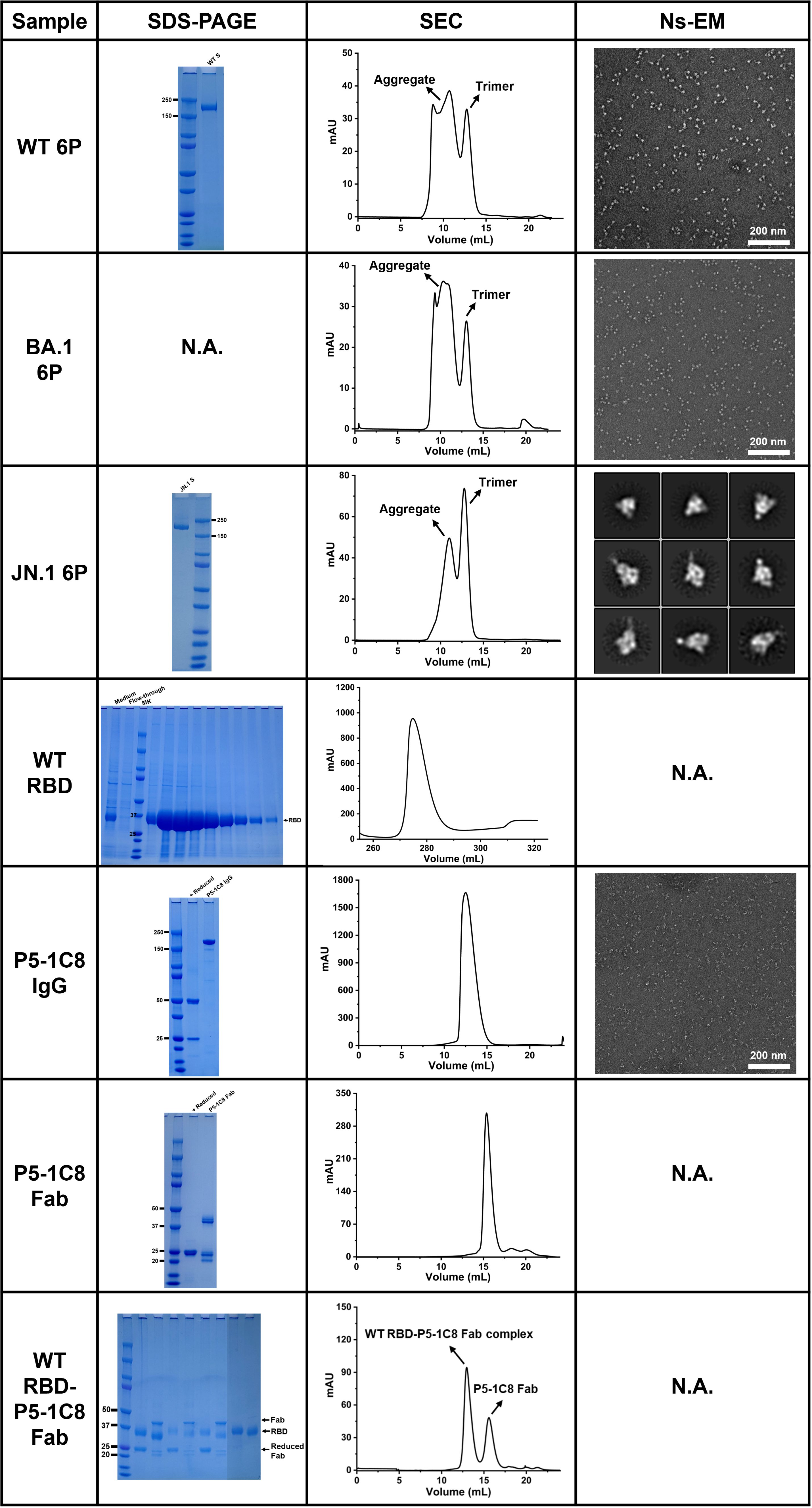

**Table S3.** Negative‑stain EM dataset summary of antibody-spike complexes and JN.1 spike-only. Entries include construct names (with EMD accession numbers), defined structural classes, particle numbers per class, raw micrographs, 2D class average, and 3D reconstruction. N.A.: not available.

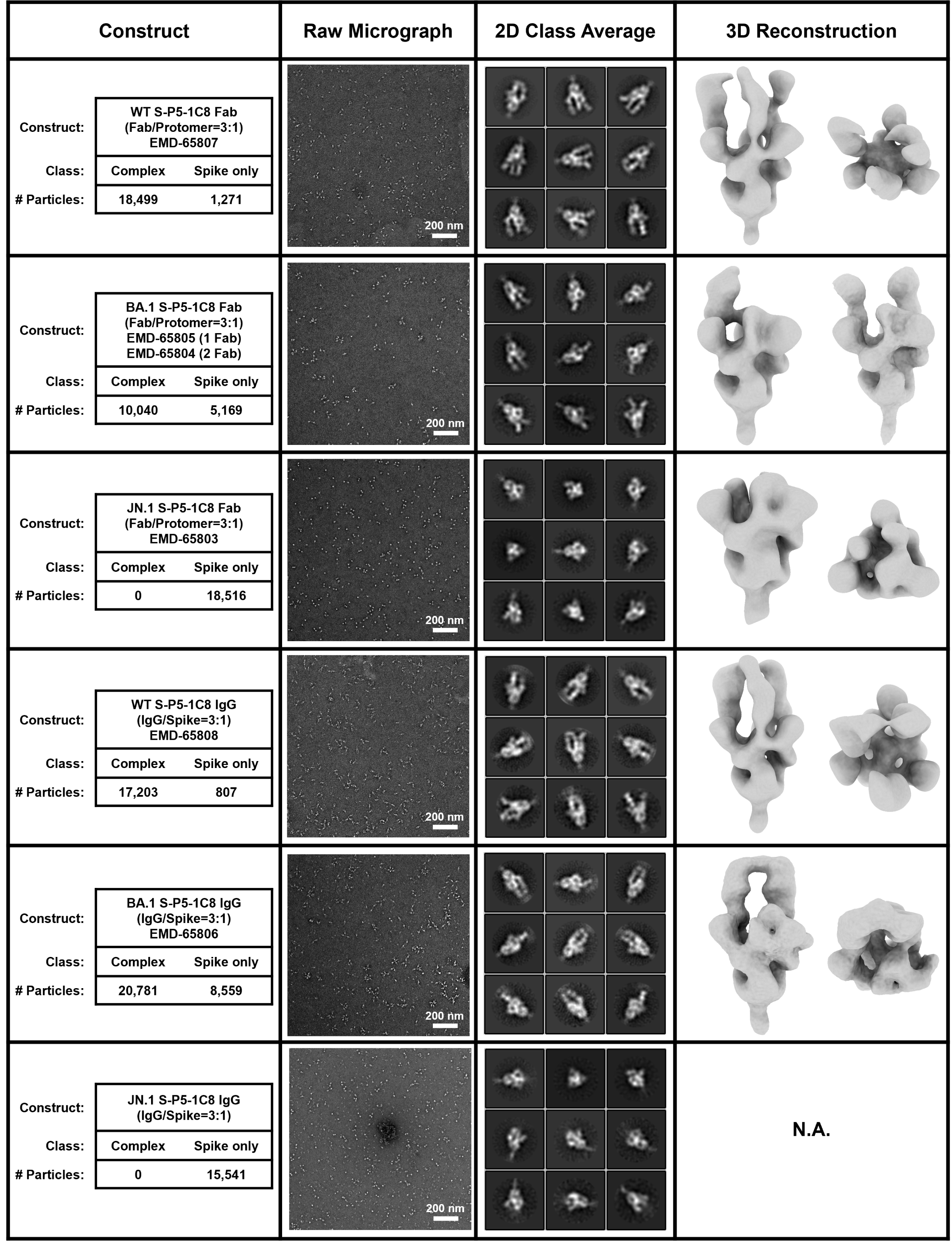

**Table S4.** The hydrogen bonds and salt bridges at the SARS-CoV-2 WT RBD and P5-1C8 Fab interfaces (distance cutoff 4 Å).

| **Hydrogen bonds** | | | |
| --- | --- | --- | --- |
| RBD | H chain | RBD | L chain |
| A475-O | I28-N | K417-NZ | Y33-OH |
| A475-O | N32-ND2 |  |  |
| L455-O | Y33-OH |  |  |
| Y421-OH | S53-N |  |  |
| R457-O | S53-OG |  |  |
| Y421-OH | G54-N |  |  |
| D420-OD2 | S56-OG |  |  |
| N487-OD1 | R97-NH1 |  |  |
| N487-OD1 | R97-NH2 |  |  |
| Y453-OH | Y102-OH |  |  |
| K417-NZ | D101-OD1 |  |  |
| K417-NZ | D101-OD2 |  |  |
| Y473-OH | R31-O |  |  |
| Q493-NE2 | Y99-OH |  |  |
| **Salt bridges** | |  |  |
| RBD | H chain |  |  |
| K417-NZ | D101-OD1 |  |  |
| K417-NZ | D101-OD2 |  |  |

Table S5. Crystallization kits are utilized during initial screening trials.

| **Swissci-Plate 96-3** | **Kit Name** | **Kit Brand** |
| --- | --- | --- |
| S-3 | Crystal Screen 1/2 | Hampton Research |
| S-4 | PEG Rx 1/2 | Hampton Research |
| S-6 | Index | Hampton Research |
| S-7 | PEG/Ion 1/2 | Hampton Research |

**Table S6.** Cryo-EM Data collection parameters of SARS-CoV-2 WT S-P5-1C8 IgG complexes.

| Sample | SARS-CoV-2 WT S-P5-1C8 IgG  EMD-65801 (1.5 IgG)  EMD-65802 (1 IgG) |
| --- | --- |
| **Data collection and processing** |  |
| Magnification | 29000x |
| Voltage (kV) | 300 |
| Electron exposure (e^-^/Å^2^) | 50 |
| Defocus range (μm) | -1 to -1.6 |
| Pixel size (Å) | 0.97 |
| Micrographs (No.) | 940 |
| Initial particles (No.) | 328,106 |
| Final particles (No.) | 152,697 |
| Map resolution (Å) | 5.48 |
| Local resolution range (Å) | 7.75 |
| Map sharpening B-factor (Å^2^) | 302.9 |

**Table S7.** Data collection and refinement statistics (molecular replacement).

|  | SARS-CoV-2 WT RBD & P5-1C8 Fab  (PDB-9K6J) |
| --- | --- |
| **Data collection** |  |
| Space group | P 21 21 21 |
| Unit cell dimensions |  |
| *a, b, c* (Å) | 55.306 115.092 144.135 |
| *α, β, γ* (°) | 90.0 90.0 90.0 |
| Resolution range (Å) | 89.94 - 2.39 (2.52 - 2.39) |
| Rmerge (%) | 0.459 (1.537) |
| *I/σ I* | 4.400 (1.500) |
| Completeness (%) | 100.0 (100.0) |
| Redundancy | 9.7 (9.0) |
| CC1/2 | 0.951 (0.413) |
| **Refinement** |  |
| Resolution (Å) | 53.44-2.39 |
| No. reflections | 34150 (3575) |
| *Rwork/Rfree* (%) | 23.52/28.18 |
| No. atoms |  |
| Protein | 4597 |
| Ligands | 14 |
| Water | 157 |
| *B-factor* (Å^2^) |  |
| Protein | 23.58 |
| Ligands | 61.27 |
| Water | 21.15 |
| R.m.s deviations |  |
| Bond length (Å) | 0.292 |
| Bond angles (°) | 0.465 |
| Ramachandran plot |  |
| Favored (%) | 96.98 |
| Allowed (%) | 3.02 |
| Outliers (%) | 0.00 |

**Table S8.** Summary of coarse-grained MD simulation parameters as deﬁned in the Supporting Information. All simulations were conducted with spike-decorated nanoparticles (NPs) bearing 8 spike trimers per NP to align with experimental conditions.

| **Category** | **Parameter** | **Value / Setting** | **Source (SI)** |
| --- | --- | --- | --- |
| Unit  System | Length unit (σ) | 7*.*5 nm | Eq. (1) |
|  | Time unit (t_0_) | 0*.*1 µs | Eq. (1) |
|  | Energy unit (ε) | *k_B_T* at 300 K = 4*.*14 × 10*^−^*^21^ J | Eq. (1) |
| Molecular  Dimensions | Nanoparticle radius (R_np_) | 7*.*5 nm | Text |
|  | Spike length (L_spike_) | 25 nm | Text |
|  | IgG length (L_IgG_) | 15 nm | Text |
| Molecular  Masses | Spike mass (M_Spike_) | 540 kg*/*mol | Text |
|  | IgG mass (M_IgG_) | 150 kg*/*mol | Text |
| Concentrations | Spike on NP | 0*.*5 µM (8 spike trimers per NP) | Text |
|  | IgG | 1*.*4 µM | Text |
| Simulation  Setup | Software | GALAMOST | Ref. [10] |
|  | Ensemble | NVT | Text |
|  | Thermostat | Velocity rescaling  (τ = 0*.*2 ps) | Text |
|  | Barostat | Nose-Hoover  (τ = 1*.*0 ps, parameter = 0.5) | Text |
|  | Time step | 4 ns | Text |
|  | Total duration | 15 µs | Text |
|  | Box size | 1500×1500×1500 nm^3^ | Text |
|  | Trajectory save interval | Every 1 ns | Text |
| Interaction  Potentials | Bonded | Harmonic bond/angle + dihedrals | Eqs.(2), (3) |
|  | Non-bonded | LJ + Coulomb (reaction ﬁeld) | Eqs.(4), (5) |
|  | Force Field | MARTINI parameterization | Text |

**Legends for Movies S1 to S2:**

**Movie S1.** Coarse-grained molecular dynamics simulation of NP-WT S nanoparticles interacting with full-length P5-1C8 IgG. The system contains 1.4 μM IgG and ~0.5 μM WT spike trimer displayed on nanoparticles. Nanoparticles are shown in grey. WT spikes are segmented by amino-acid numbering into four domains—residues 1-527 (NTD and RBD), 528-833 (CTD1, CTD2, and FP), 834-1162 (HR1, CH, and CD), and 1163-1240 (HR2 and foldon)—and rendered as a continues blue gradient from top (intense blue) to bottom (faded blue). P5-1C8 IgG is shown in pink, with both Fab domains in pink and the hinge region and Fc domain in pale pink.

**Movie S2.** Coarse-grained molecular dynamics simulation of NP-JN.1 S nanoparticles interacting with full-length P5-1C8 IgG. The system contains 1.4 μM IgG and ~0.5 μM JN.1 spike trimer displayed on nanoparticles. Nanoparticles are shown in grey. JN.1 spike are segmented by amino-acid numbering into four domains—residues 1-527 (NTD and RBD), 528-833 (CTD1, CTD2, and FP), 834-1162 (HR1, CH, and CD), and 1163-1240 (HR2 and foldon)—and rendered as a continues green gradient from top (intense green) to bottom (faded green). P5-1C8 IgG is shown in pink: both Fab domains are pink while the hinge region and Fc domain are rendered in pale pink.
