## Supplementary material for "IgG-Bridging–Seeded Synergistic Aggregation of SARS-CoV-2 Spikes Underlies Potent Neutralization by A Low-Affinity Antibody": Validation Report

### Full wwPDB EM Validation Report (i)

Oct 10, 2025 – 08:31 PM JST

| | | |
|---|---|---|
| EMDB ID | : | EMD-65807 |
| Title | : | Immune complex of P5-1C8 Fab binding the RBD of SARS-CoV-2 WT 6p spike protein |
| Deposited on | : | 2025-08-11 |
| Resolution | : | 25.00 Å(reported) |

**This wwPDB validation report is for manuscript review**

This is a Full wwPDB EM Validation Report.

This report is produced by the wwPDB biocuration pipeline after annotation of the structure.

We welcome your comments at **
A user guide is available at
https://www.wwpdb.org/validation/2017/EMMapValidationReportHelp
with specific help available everywhere you see the (i) symbol.

The types of validation reports are described at
http://www.wwpdb.org/validation/2017/FAQs#types.

For Manuscript Review

The following versions of software and data (see references (i)) were used in the production of this report:

| | | |
|---|---|---|
| EMDB validation analysis | : | 0.0.1.dev129 |
| Validation Pipeline (wwPDB-VP) | : | 2.46 |

### 1 Experimental information (i)

| Property | Value | Source |
|---|---|---|
| EM reconstruction method | SINGLE PARTICLE | Depositor |
| Imposed symmetry | POINT, Not provided | |
| Number of particles used | 5775 | Depositor |
| Resolution determination method | FSC 0.5 CUT-OFF | Depositor |
| CTF correction method | PHASE FLIPPING AND AMPLITUDE CORRECTION | Depositor |
| Microscope | JEOL 2100F | Depositor |
| Voltage (kV) | 200 | Depositor |
| Electron dose ($e^-/\text{Å}^2$) | 25 | Depositor |
| Minimum defocus (nm) | 1500 | Depositor |
| Maximum defocus (nm) | 1500 | Depositor |
| Magnification | Not provided | |
| Image detector | OTHER | Depositor |
| Maximum map value | 0.090 | Depositor |
| Minimum map value | -0.033 | Depositor |
| Average map value | 0.000 | Depositor |
| Map value standard deviation | 0.005 | Depositor |
| Recommended contour level | 0.027 | Depositor |
| Map size (Å) | 424.32, 424.32, 424.32 | wwPDB |
| Map dimensions | 192, 192, 192 | wwPDB |
| Map angles (°) | 90.0, 90.0, 90.0 | wwPDB |
| Pixel spacing (Å) | 2.21, 2.21, 2.21 | Depositor |

For Manuscript Review

### 2 Map visualisation (i)

This section contains visualisations of the EMDB entry EMD-65807. These allow visual inspection of the internal detail of the map and identification of artifacts.

Images derived from a raw map, generated by summing the deposited half-maps, are presented below the corresponding image components of the primary map to allow further visual inspection and comparison with those of the primary map.

#### 2.1 Orthogonal projections (i)

##### 2.1.1 Primary map

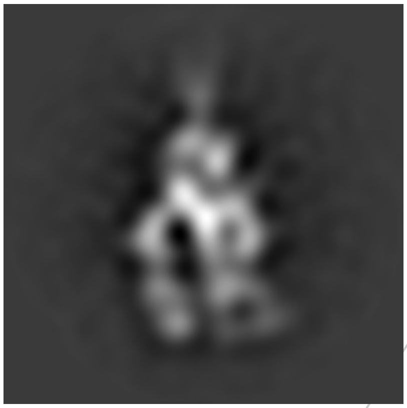
X

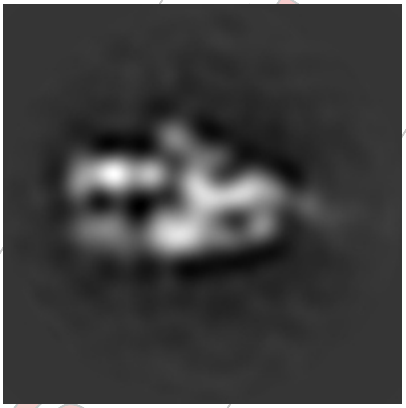
Y

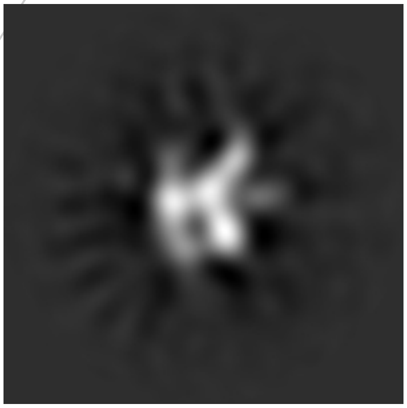
Z

##### 2.1.2 Raw map

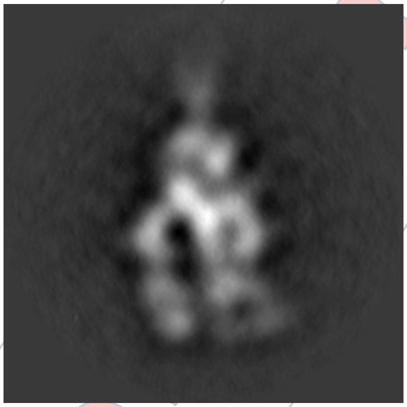
X

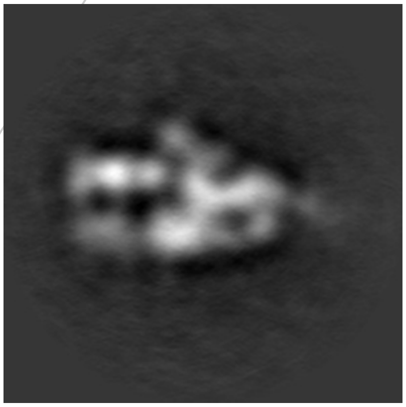
Y

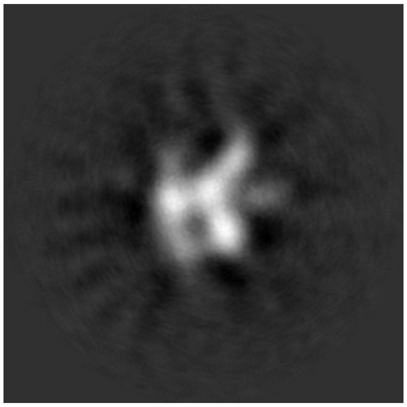
Z

The images above show the map projected in three orthogonal directions.

For Manuscript Review

#### 2.2 Central slices (i)

##### 2.2.1 Primary map

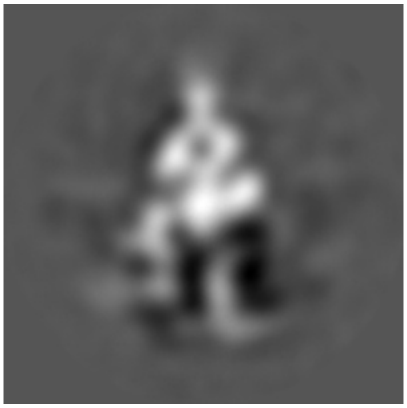

X Index: 96

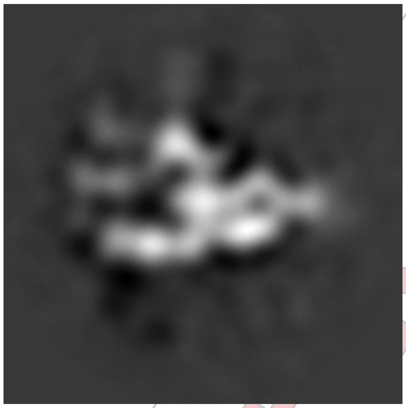

Y Index: 96

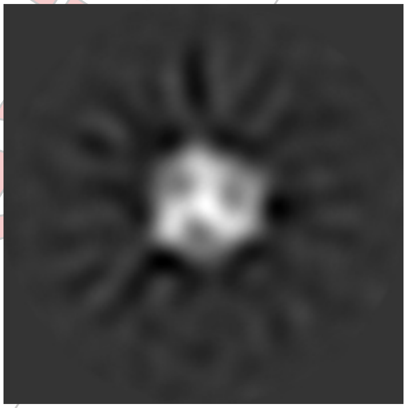

Z Index: 96

##### 2.2.2 Raw map

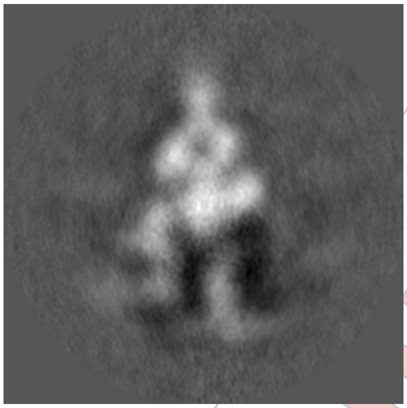

X Index: 96

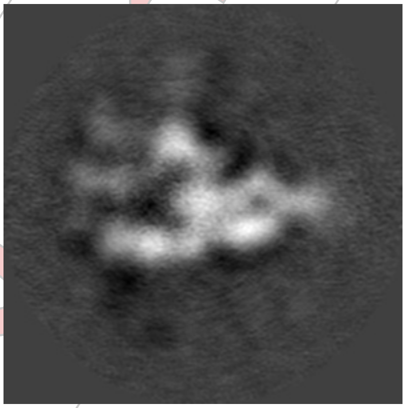

Y Index: 96

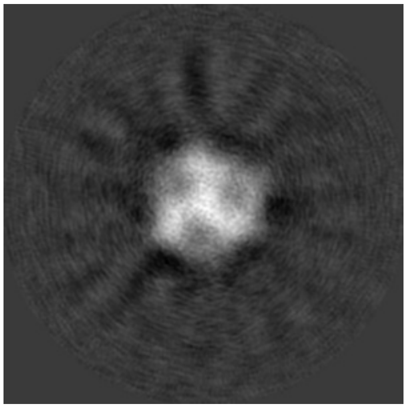

Z Index: 96

The images above show central slices of the map in three orthogonal directions.

For Manuscript Review

#### 2.3 Largest variance slices (i)

##### 2.3.1 Primary map

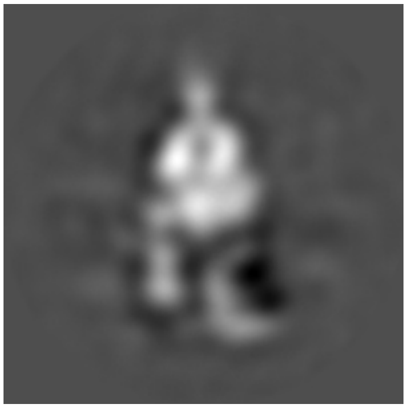
X Index: 100

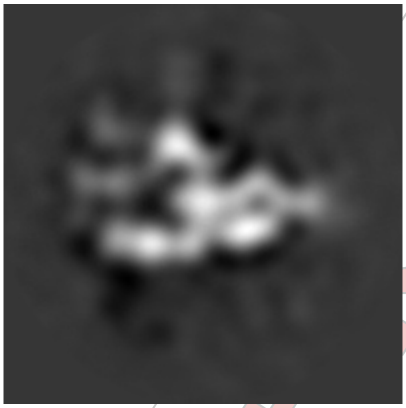
Y Index: 97

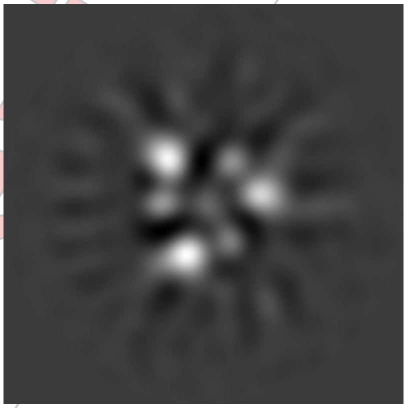
Z Index: 81

##### 2.3.2 Raw map

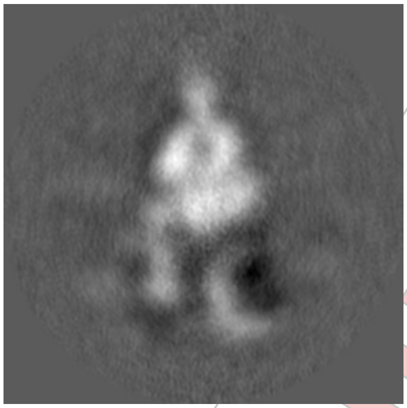
X Index: 99

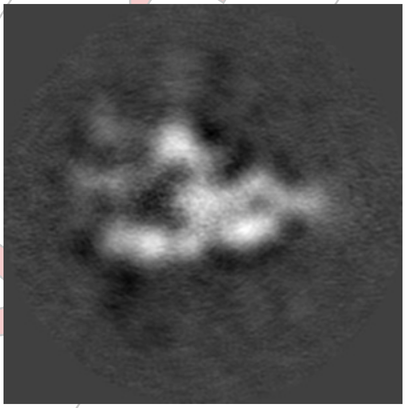
Y Index: 97

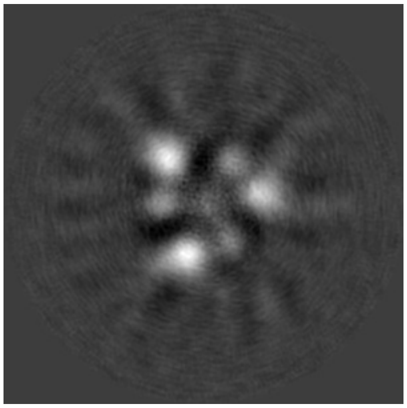
Z Index: 81

The images above show the largest variance slices of the map in three orthogonal directions.

For Manuscript Review

#### 2.4 Orthogonal standard-deviation projections (False-color) (i)

##### 2.4.1 Primary map

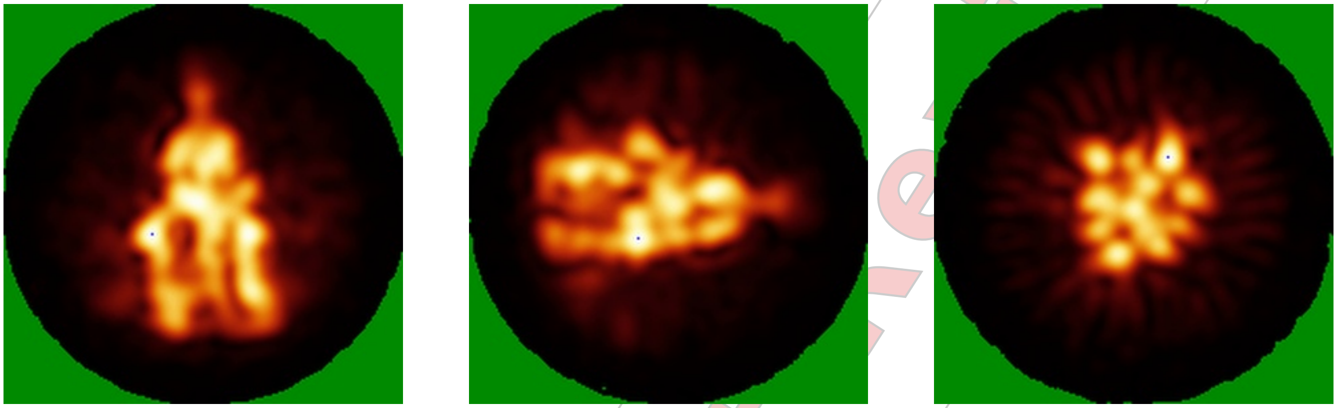

X Y Z

##### 2.4.2 Raw map

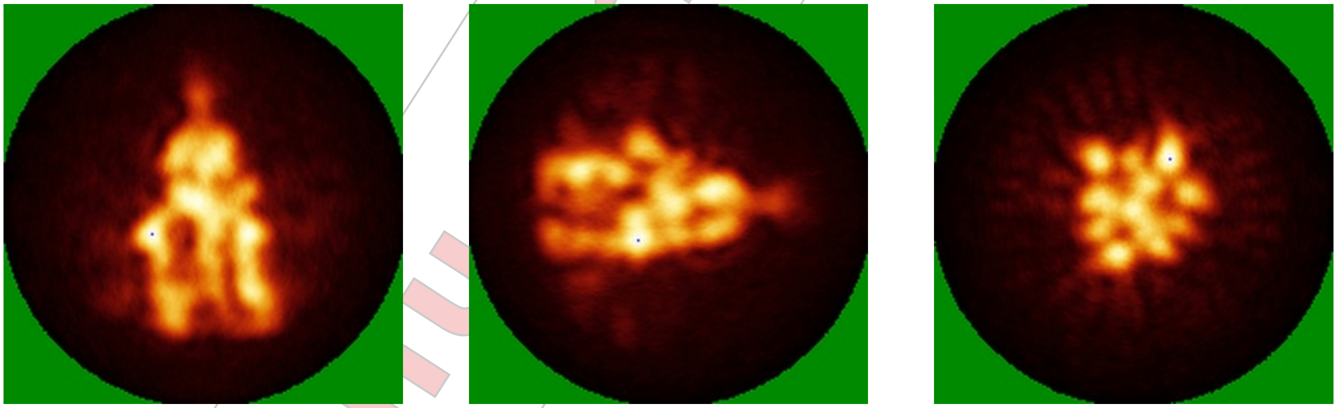

X Y Z

The images above show the map standard deviation projections with false color in three orthogonal directions. Minimum values are shown in green, max in blue, and dark to light orange shades represent small to large values respectively.

For Manuscript Review

#### 2.5 Orthogonal surface views

##### 2.5.1 Primary map

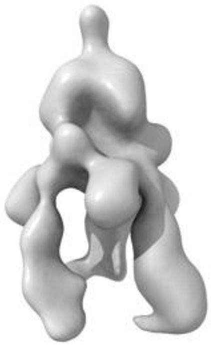

X

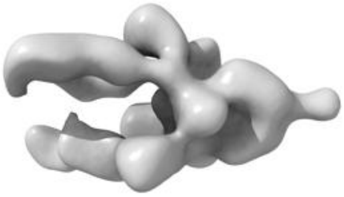

Y

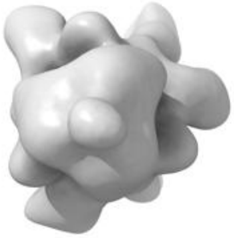

Z

The images above show the 3D surface view of the map at the recommended contour level 0.027. These images, in conjunction with the slice images, may facilitate assessment of whether an appropriate contour level has been provided.

##### 2.5.2 Raw map

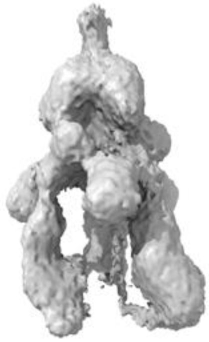

X

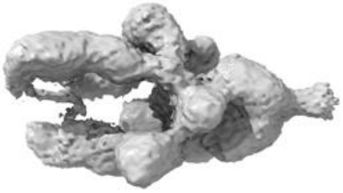

Y

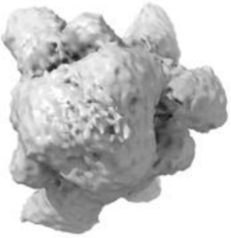

Z

These images show the 3D surface of the raw map. The raw map's contour level was selected so that its surface encloses the same volume as the primary map does at its recommended contour level.

#### 2.6 Mask visualisation

This section was not generated. No masks/segmentation were deposited.

For Manuscript Review

### 3 Map analysis (i)

This section contains the results of statistical analysis of the map.

#### 3.1 Map-value distribution (i)

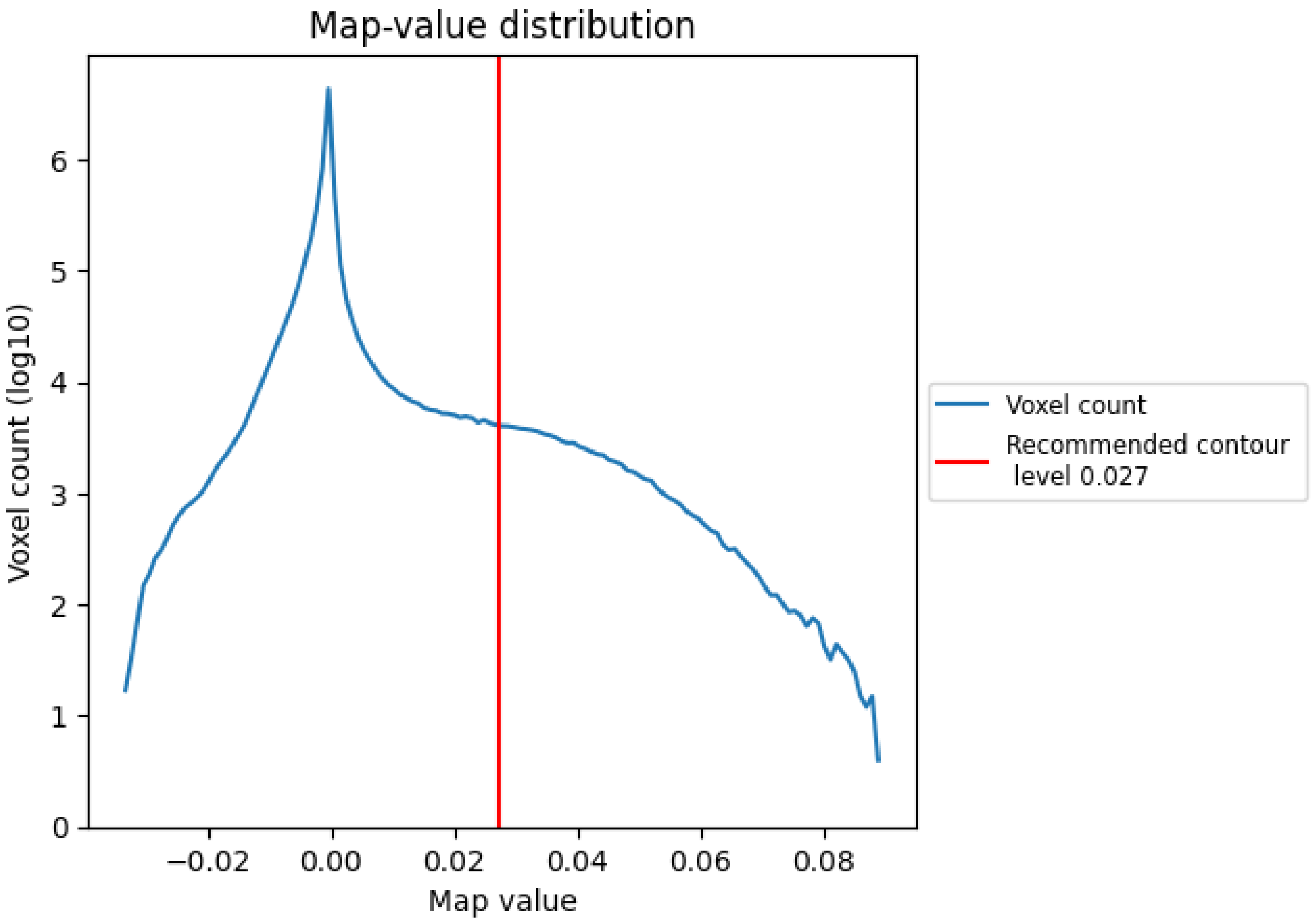


The map-value distribution is plotted in 128 intervals along the x-axis. The y-axis is logarithmic. A spike in this graph at zero usually indicates that the volume has been masked.

For Manuscript Review

#### 3.2 Volume estimate ⓘ

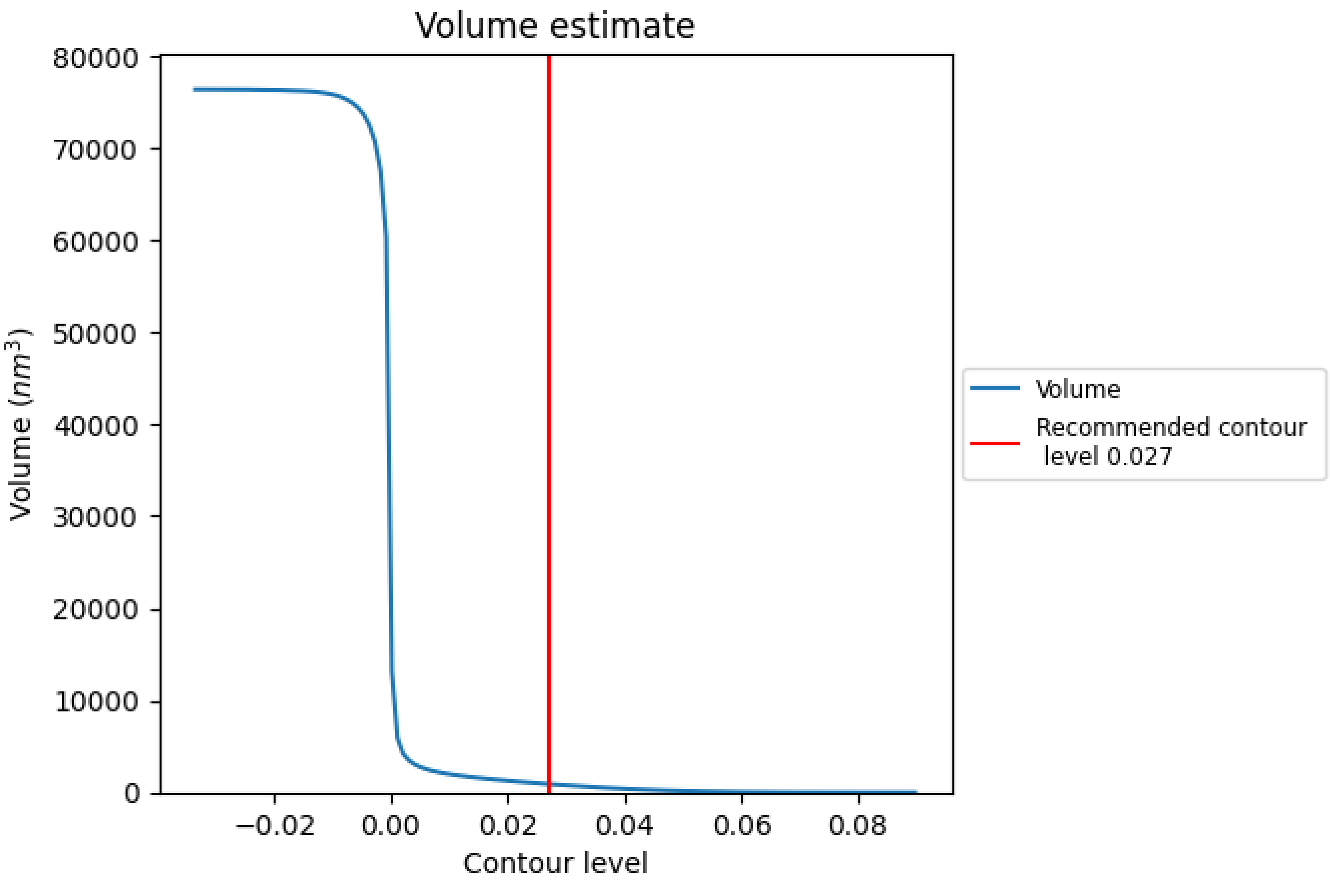


The volume at the recommended contour level is 913 $nm^3$; this corresponds to an approximate mass of 825 kDa.

The volume estimate graph shows how the enclosed volume varies with the contour level. The recommended contour level is shown as a vertical line and the intersection between the line and the curve gives the volume of the enclosed surface at the given level.

For Manuscript Review

#### 3.3 Rotationally averaged power spectrum (i)

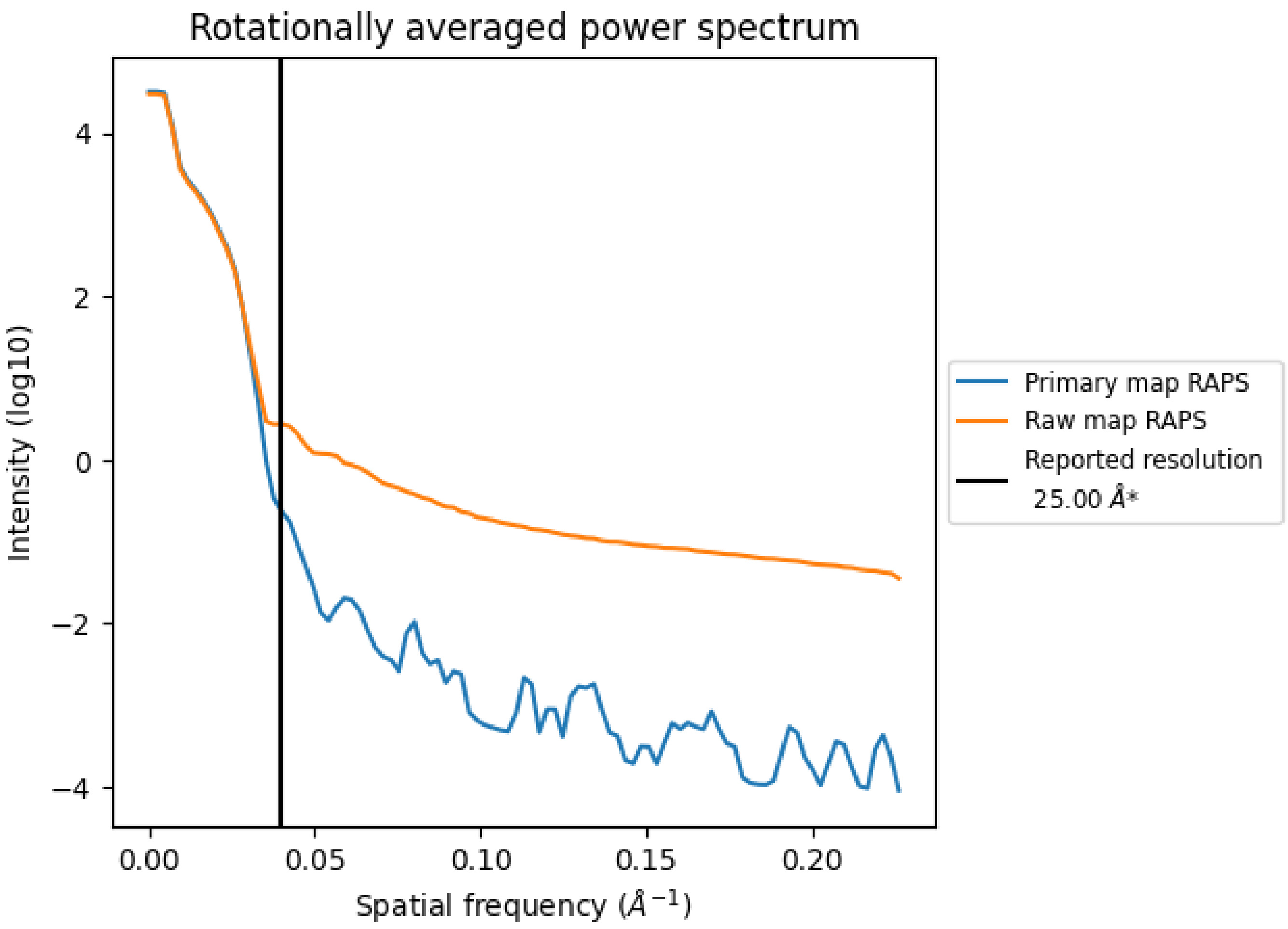


*Reported resolution corresponds to spatial frequency of 0.040 Å$^{-1}$

For Manuscript Review

### 4 Fourier-Shell correlation ⓘ

Fourier-Shell Correlation (FSC) is the most commonly used method to estimate the resolution of single-particle and subtomogram-averaged maps. The shape of the curve depends on the imposed symmetry, mask and whether or not the two 3D reconstructions used were processed from a common reference. The reported resolution is shown as a black line. A curve is displayed for the half-bit criterion in addition to lines showing the 0.143 gold standard cut-off and 0.5 cut-off.

#### 4.1 FSC ⓘ

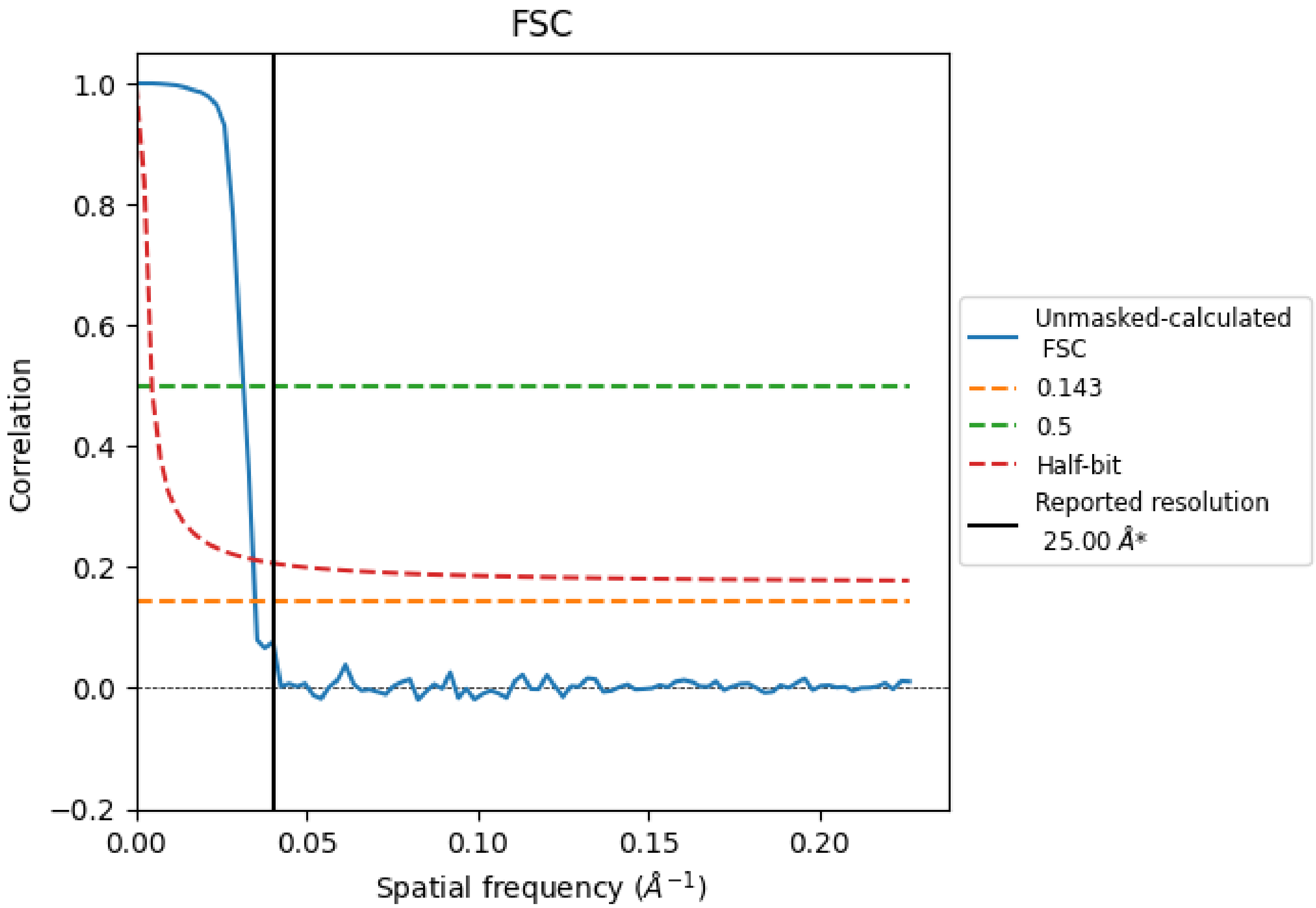


*Reported resolution corresponds to spatial frequency of 0.040 Å$^{-1}$

For Manuscript Review

#### 4.2 Resolution estimates (i)

| Resolution estimate (Å) | Estimation criterion (FSC cut-off) | | |
|---|---|---|---|
| | 0.143 | 0.5 | Half-bit |
| Reported by author | - | 25.00 | - |
| Author-provided FSC curve | - | - | - |
| Unmasked-calculated* | 28.74 | 31.85 | 29.24 |

*Resolution estimate based on FSC curve calculated by comparison of deposited half-maps. The value from deposited half-maps intersecting FSC 0.5 CUT-OFF 31.85 differs from the reported value 25.0 by more than 10 %

For Manuscript Review
